# Transcriptomic analysis revealed reactive oxygen species scavenging mechanisms associated with ferrous iron toxicity in aromatic Keteki Joha rice

**DOI:** 10.1101/2021.10.13.464192

**Authors:** Preetom Regon, Sangita Dey, Mehzabin Rehman, Amit Kumar Pradhan, Bhaben Tanti, Anupam Das Talukdar, Sanjib Kumar Panda

**Affiliations:** Department of Life Science and Bioinformatics, Assam University, Silchar-788011, Assam, India; Department of Botany, Gauhati University, Guwahati-781014, Assam, India; Department of Biochemistry, Central University of Rajasthan, Ajmer- 305817, India

**Keywords:** Fe^2+^ toxicity, RNA-Seq, Transcriptome, Fe homeostasis, ROS

## Abstract

Lowland acidic soils with water-logged regions are often affected by ferrous iron (Fe^2+^) toxicity, a major yield-limiting factor of rice production. The Reactive Oxygen Species (ROS) was hypothesized to be crucial under severe Fe^2+^ toxicity conditions. However, molecular mechanisms and associated ROS homeostasis genes are still not well-explored. In this study, a comparative RNA-Seq based transcriptome analysis was conducted to understand the Fe^2+^ toxicity tolerance mechanism in aromatic Keteki Joha. About 69 Fe homeostasis related genes and their homologs were identified, where most of the genes were downregulated. Differentially expressed genes (DEGs) are associated with biological processes-response to stress, stimulus and abiotic stimulus. DEGs involved in the Biosynthesis of amino acids, RNA degradation, Glutathione metabolism etc. were induced, whereas, Phenylpropanoid biosynthesis, Photosynthesis, and Fatty acid elongation were inhibited. The *Mitochondrial iron transporter* (*OsMIT*), *Vacuolar Iron Transporter 2* (*OsVIT2*), *Ferritin* (*OsFER*), *Vacuolar Mugineic Acid Transporter* (*OsVMT*), *Phenolic Efflux Zero1* (*OsPEZ1*), *Root Meander Curling* (*OsRMC*), *Nicotianamine synthase* (*OsNAS3*), etc. were upregulated in different tissues suggesting the importance of Fe retention and sequestration for detoxification. However, several antioxidants, ROS scavenging genes and abiotic stress-responsive transcription factors indicate ROS homeostasis as one of the most important defense mechanisms under severe Fe^2+^ toxicity. The CAT, GSH, APX, MDHAR, DHAR, GR were upregulated. Moreover, abiotic stress-responsive transcription factors NAC, MYB, ARF, bZIP, WRKY, C2H2-ZFP were also upregulated. Accordingly, ROS homeostasis has been proposed to be an important defense mechanism under such conditions. Thus, our results will enrich the knowledge of understanding Fe-homeostasis in rice.

## 1. Introduction

Iron (Fe) is an essential micronutrient participating in various vital processes in plants due to its redox status between ferrous (Fe^2+^) and ferric (Fe^3+^) forms. Fe functions as an electron acceptor or donor actively participating in photosynthesis and respiration ^1,2^. Lowland and flooded soil are poor in oxygen concentration and become acidic. The flooding of rice fields leads to the reduction of Fe^3+^ to Fe^2+^, which becomes highly toxic to rice. Fe^2+^ toxicity is a major cause of abiotic stress affecting large rice-growing areas, especially in Asian countries ^3^. Depending on stress intensities, a loss of 12% to 100% has been reported on rice yields ^4,5^. Fe^2+^ toxicity is a severe agricultural constrain that generally occurs in acidic soils. Thus, understanding the underlying molecular mechanisms associated with their adaptation will help us to build strategies for future rice breeding programmes for Fe^2+^ toxicity tolerance.

Higher plants have evolved two acquisition strategies to overcome the limited Fe availability *viz*., strategy I and strategy II. Strategy I is based on the acidification of rhizospheres by releasing protons from the roots, which reduces Fe^3+^ to Fe^2+^ and its transportation through iron-regulated transporter (IRT) across the root plasma membrane. Strategy II is based on the uptake of Fe^3+^-phytosiderophore complexes. Although being a Strategy II plant, rice *(Oryza sativa* L.) possesses Fe^2+^ transporter genes (*Os*IRT1 and *Os*IRT2) that can directly uptake Fe^2+^ from the soil ^6,7^. The mechanism of Fe^2+^ toxicity tolerance in rice plants has been highlighted in several studies ^8–12^ Rice plants possess several defense mechanisms to cope with Fe^2+^ toxicity. The defense mechanism of rice plants for Fe^2+^ tolerance can be divided into four mechanisms ^12^. Defense 1 (Fe Exclusion from Roots) is a root-based tolerance mechanism where Fe plaques (Fe^3+^ precipitation) on the root surface acts as a barrier to the uptake of Fe^2+^ into root tissues. Fe plaque is formed due to the rhizospheric oxidation of Fe^2+^ to Fe^3+^ by oxygen transport from shoots to roots ^3,8^. Defense 2 (Fe Retention in Roots and Suppression of Fe Translocation to Shoots) where discrimination center and *OsFER1* plays an important role by retaining excess Fe in root and avoid translocation into the shoot. Defense 3 (Fe Compartmentalization in Shoots) includes compartmentalization, disposal, or storage of Fe inside shoots by *OsVIT1*, *OsVIT2*, and *OsFER1/2*. Defense 4 (ROS Detoxification) includes enzymatic detoxification, scavenging of ROS by antioxidants. Defense 1-3 was highlighted to be work on mild to moderate Fe excess conditions and do not seriously affect rice growth. In contrast, under Fe severe conditions, defense 4 was hypothesized to be work at a molecular level that causes bronzing and inhibits plant growth and development ^12^. ROS antioxidants like glutathione-S-transferase and ascorbate oxidase were reported to play an important role in shoot-based Fe tolerance in rice plants ^10^. Besides, under severe Fe excess conditions, cytochrome P450 family proteins, *OsNAC4*, *OsNAC5*, and *OsNAC6*, played an important role in alleviating ROS ^12^. The elucidation of the genes involved in response to Fe-toxicity is fundamental to understanding the mechanism that confers tolerance to stress and the development of tolerant cultivars. With the advancement of NGS technology, the RNA-seq technique has become a valuable tool for transcriptional profile analysis, providing a better understanding of gene responses to Fe^2+^ toxicity. To date, many studies on the transcriptional profile of rice under Fe-stress have been done. However, there is still a broad spectrum that needs better clarification to understand the Fe stress mechanism, which can help to understand the unsolved queries related to Fe toxicity in rice.

In the present study, the RNA-seq technique was used to analyze the transcriptional profile of aromatic Keteki Joha under severe Fe^2+^ toxic conditions to elucidate the differentially expressed genes. In addition, the gene ontology (GO) and Kyoto Encyclopedia of Genes and Genomes (KEGG) were used to demonstrate major metabolic pathways affected in response to Fe^2+^ toxicity. This investigation may provide important transcripts information that will further elucidate the Fe homeostasis gene in Fe^2+^ toxicity and scope to investigate the possible adaptation mechanism in rice plants.

## 2. Materials and Methods

### 2.1 Plant Growth and Treatment

The seeds of Keteki Joha were collected from Regional Agricultural Research Station, Titabor, Assam, India. Rice seedlings were grown under controlled growth conditions, and after three weeks, rice seedlings were treated with 2.5 mM of FeEDTA for 72 h, as described previously ^13^. The seedlings were treated in Hoagland hydroponic solution, whereas the mature stage was treated in soil. For the mature stage, rice plants were grown in a pot containing approximately 3.5 kg of soil (dry weight). About 60-70-day old plants were treated with 5 L of 2.5 mM FeEDTA for two weeks, and leave samples were harvested for RNA isolation and sequencing (Fig S1). The FeEDTA solution was changed every 96 h. Three utterly independent pot experiments were considered as biological replicates (Fig S2).

### 2.2 RNA-Sequencing

RNA was isolated from both roots and leaves with 3 biological replicates using the RNAqueous Phenol-free total RNA isolation kit of Invitrogen, Thermo Fisher Scientific. RNA Integrity Number (RIN) value of more than 6.5 was used to prepare the sequencing library prepared by Truseq Stranded RNA Library Prep Kit-Plant. In addition, 100-150 bp RNA Seq was done through Illumina HiSeq 2000 platform.

### 2.3 Data-processing, Assembly and differential expression

Basic statistics of the sequencing raw reads were accessed by FastQC (https://www.bioinformatics.babraham.ac.uk/projects/fastqc/). Raw reads were trimmed with Trimmomatic to remove sequencing adapters ^14^. Reference-based assembly was performed by STAR, where Samtools was used to manipulate the bam files ^15,16^. Rice Genome Annotation Project (http://rice.plantbiology.msu.edu/) was used as a reference sequence database for assembly ^17^. Transcript abundance was quantified using the featureCounts program of the Subread package 18. Differential gene expression was performed using the DESeq2 package of the R program ^19,20^. EnhancedVolcano and ComplexHeatmap package of R was used to represent the volcano and heatmap plots of the DEGs ^21,22^.

### 2.4 Annotation

Transcripts were annotated using an in-house pipeline. Briefly, rice annotation data available in the UniProt database were downloaded, and local BLASTX was performed ^23,24^. The best scoring results were merged with the transcript using the R program ^19^. Gene Ontology, Sub-cellular localization, cross reference KEGG ID, and Pfam were considered during the UniProt annotation. Additional gene description was assigned using the gene annotation data available in the Rice Genome Annotation Project (http://rice.plantbiology.msu.edu/) ^17^. Also, the corresponding RAPD gene id and gene symbol were assigned to the annotated transcript by using the R program ^25^.

### 2.5 Gene set enrichment analysis

Gene set enrichment analysis was done with the topGO of the R package ^26^. The Gene ontology term of each locus was obtained from Rice Genome Annotation Project (http://rice.plantbiology.msu.edu/), and Molecular function, biological process and cellular component were studied for the selected top 1000 DEGs (500 Up/ 500 down) ^17^. Gene ontology terms were ranked by employing the Fishers’ exact statistical test and classic algorithm of topGO. The ggplot2 package of R program was used to represent the graphical representation ^27^.

Metabolic pathways were studied using the Kyoto Encyclopedia of Genes and Genomes (KEGG) database ^28^. Briefly, all the protein sequences of the DEGs were downloaded from the Rice Genome Annotation Project database and reannotated using GhostKOALA in the KEGG database ^29^. Cross-reference KEGG gene ID of *Oryza sativa*, Japonica (osa), KEGG entry no T01015, and GHOSTX score >=100 were considered for pathway mapping. Finally, the KEGG pathway was analyzed using the KEGG mapper. Expression value (log2FoldChange) of the DEGs were integrated into their corresponding KEGG gene of each sample, and multiple states pathway maps were rendered by the Pathview web tool ^30^.

### 2.6 Differential exon-usage of the DEGs

Differential exon usage of the DEGs was inferred by Bioconductor package DEXSeq ^31^. Briefly, the mapping was done with STAR and transcript abundance was quantified with featureCounts as described above. The Subread to DEXSeq python script (https://github.com/vivekbhr/SubreadtoDEXSeq) was used to manipulate the featureCounts data to create the DEXSeq object.

### 2.7 Data Availability

The raw data of RNA-seq used in the study have been deposited in the Sequence Read Archive (SRA), National Center for Biotechnology Information (NCBI) under the BioProject accession number PRJNA680441 and BioSamples accessions SAMN16879912, SAMN16879913, SAMN16879914, SAMN16879915, SAMN16879916, SAMN16879917.

### 2.8 Quantitative real-time PCR analysis

The same experimental condition was maintained for quantitative real-time PCR analysis. Briefly, about 100 mg of fresh tissues with 3 biological replicates were grounded to powder with Liquid Nitrogen (LN_2_). Total RNA was isolated using the RNAqueous Phenol-free total RNA isolation kit of Invitrogen, Thermo Fisher Scientific. The cDNA synthesis was performed using the iScript Reverse Transcription Supermix for RT-qPCR kit, Bio-Rad. Gene-specific primers were designed with Primer3web (http://primer3.ut.ee/) (Table S1) and a few more were also obtained from research articles ^11,32,33^. The qRT-PCR was performed using the PowerUp SYBR Green Master Mix, Applied Biosystems and performed in QuantStudio 5 Real-Time PCR System, Applied Biosystems. Gene expression was calculated by the comparative CT method ^34^.

## 3. Results

### 3.1 RNA sequencing and reference-based transcriptome assembly

Growth of the rice plant was significantly inhibited under severe Fe^2+^ toxicity (Fig S1). RNA-Seq transcriptome was performed and approximately 09.36 to 37.40 million 100 bp sequence reads were obtained from each sequencing run from which 41.59 to 72.27% were uniquely mapped into the Japonica rice genome (Table S2 and S3). About 44% of GC content was observed among the sequencing reads. About 7708 DEGs in roots, 8271 in leaves seedling and 12885 in leaves mature stage were identified (Fig 1a).

**Figure 1:**
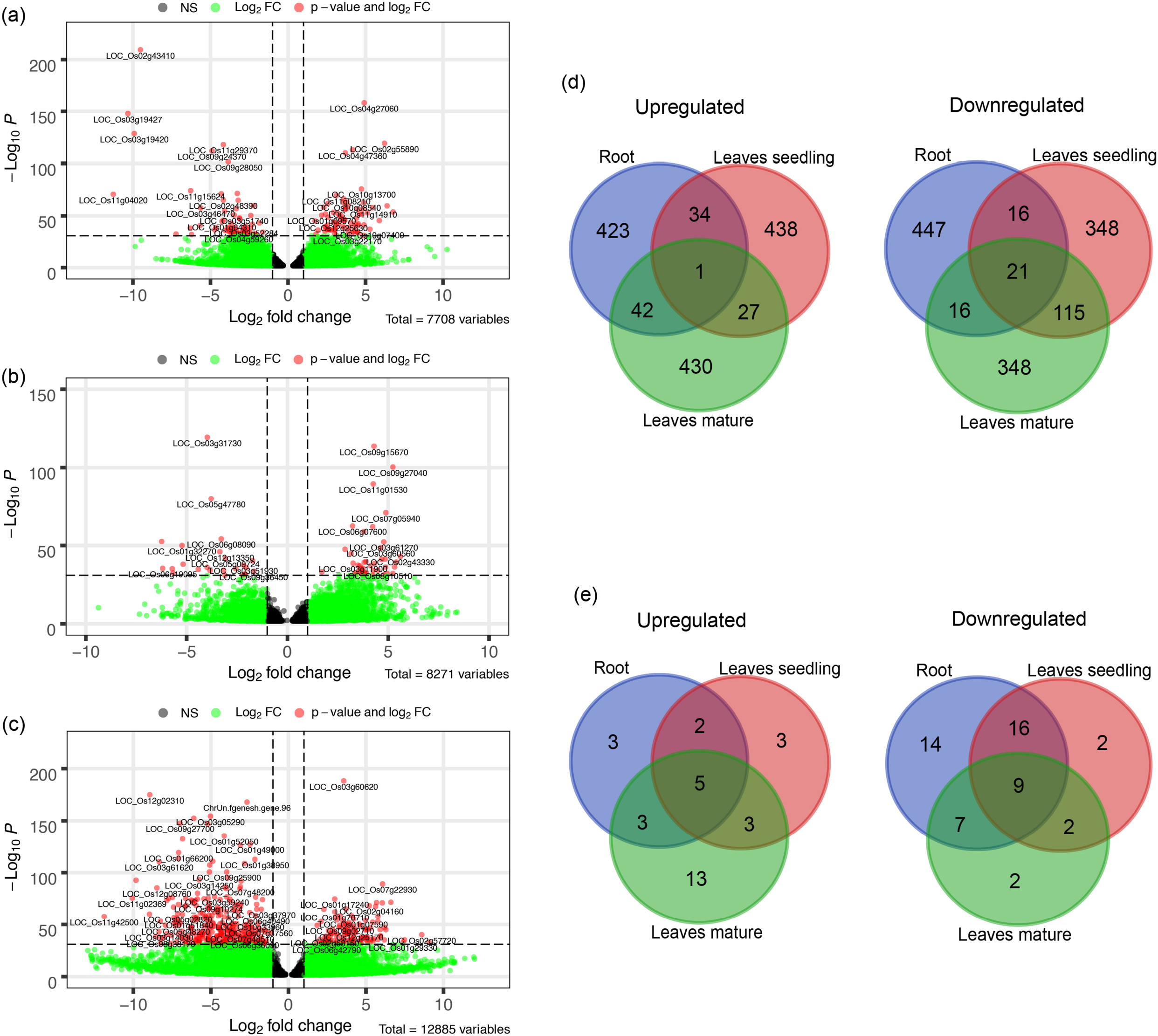
Differentially Expressed Genes under Fe^2+^ toxicity. (a-c) Volcano plots showing the expression of DEGs under different tissue samples. The p-value less than 10e-32 were tagged with their gene ID (d) Venn diagram showing the top 500 Up and downregulated common DEGs among the tissue samples (e) Venn diagram of common Fe-homeostasis related DEGs, expressed in different tissues samples. The Venn diagrams were created by using the Venn diagram software of VIB/UGent Bioinformatics & Evolutionary Genomics.

### 3.2 Differentially expressed genes in roots under Fe^2+^ toxicity

Among the DEGs, a total of 69 Fe homeostasis-related genes and their homologs were identified (Fig 2a). Most of the genes associated with Fe homeostasis were downregulated under Fe^2+^ toxicity. However, 13 Fe homeostasis related genes were upregulated in roots, of which the genes *OsPEZ1*, *OsVMT* and *OsIDI1L*/*OsARD1* were highly upregulated. Other upregulated genes include *OsNAS3*, *OsTOM3*, *OsMIT*, *OsRMC*, *OsFER1*/*OsFER2*, *OsATM3*, *OsYSL13*, *OsFRDL1*, *OsNRAMP2* and *OsbHLH58*/*OsPRI2*. The Fe^2+^ transporter *OsIRT1*, *OsIRT2*, *OsNRAMP1* and Fe/Mn/Cd transporter *OsNRAMP5* were downregulated. In addition, Fe-phytosiderophore transporters like *OsYSL15*, *OsYSL2* were also downregulated. Other downregulated genes include *OsTOM1*, *OsNAS1*, *OsNAS2*, *OsMIR*, *OsIMA1*, *OsPRPPS*, *OsDEP*, *OsIDI4*, *OsbHLH133*, *OsDMAS1*, *OsRPI*, *OsOPT7*, *OsFDH*, *OsENA1*, *OsIRO2*, *OsNRAMP1*, *OsIDI2*, *OsbHLH156*/*OsFIT* etc. (Fig 2a). Besides, few Fe homeostasis related genes like *OsVIT2*, *OsYSL10*, *OsRab6a*, *OsYSL9*, *OsTOM2*, *OsIDEF2*, *OsYSL5*, *OsPEZ2*, *OsFRO1* and *OsFRO2* were not expressed in roots (Fig 2a).

**Figure 2:**
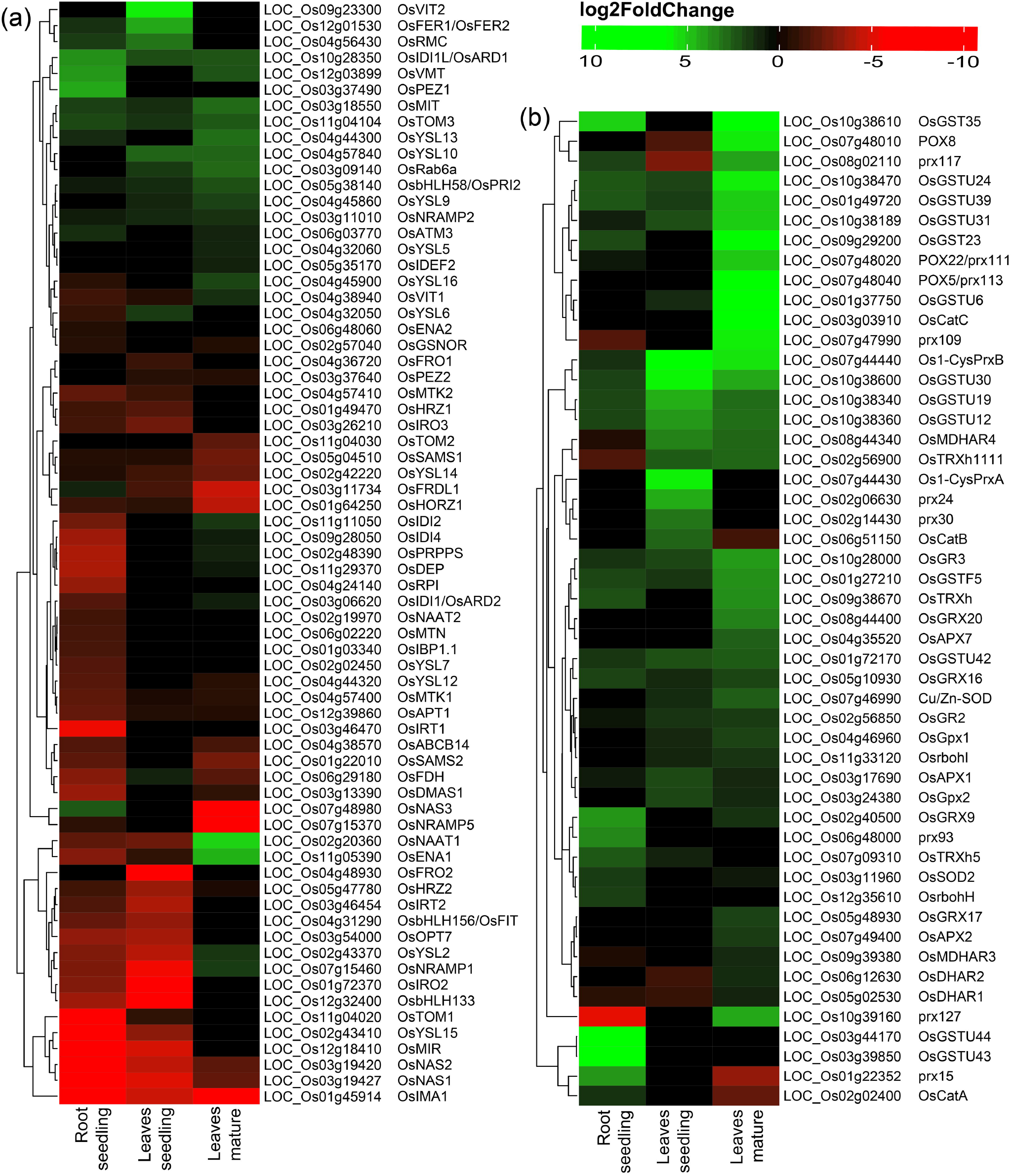
Heatmap of DEGs under Fe^2+^ toxicity. (a) Heatmap of Fe-homeostasis related genes. (b) Heatmap of antioxidant and ROS scavenging related genes. The color scale bar represents the log2FoldChange value of the DEGs.

### 3.3 Differentially expressed genes in leaves under Fe^2+^ toxicity

Variation was observed among the DEGs in the leaves of seedlings with that of the mature stage. *OsVIT2* and *OsFER1/OsFER2* were highly upregulated in the seedling stage, whereas they were not differentially expressed in the mature stage (Fig 2a). Other upregulated Fe homeostasis related genes in seedling include the *OsRMC*, *OsYSL10*, *OsIDI1L*/*OsARD1*, *OsYSL6*, *OsRab6a*, *OsTOM3*, *OsMIT*, *OsbHLH58*/*OsPRI2*, *OsNRAMP2*, *OsYSL9* and *OsFDH*. In contrast, downregulated genes include *OsFRO2*, *OsIRO2*, *OsbHLH133*, *OsNRAMP1*, *OsNAS1*, *OsMIR*, *OsIMA1*, *OsNAS2*, *OsYSL2*, *OsIRT2* etc. (Fig 2a). Among the 69 identified Fe homeostasis-related genes, 27 genes were not expressed in leaves’ seedlings (Fig 2a).

In the mature stage, genes like *OsNAAT1*, *OsENA1* were highly upregulated, followed by *OsYSL13*, *OsMIT*, *OsRab6a*, *OsYSL10*, *OsTOM3*, *OsIDI1L*/*OsARD1*, *OsVMT*, *OsbHLH58*/*OsPRI2*, etc. (Fig 2a). Most downregulated genes include *OsNAS3*, *OsIMA1*, *OsNRAMP5*, *OsFRDL1*, *OsHORZ1*, etc. (Fig 2a). A total of 25 identified Fe homeostasis related genes were differentially expressed in leaves during the mature stage (Fig 2a).

Comparative analysis of the top 500 up/down-regulated genes indicates that genes are distinctly expressed in both roots and leaves. Only one gene was found to be commonly upregulated in the roots and leaves of both seedlings as well as of the mature stage. Similarly, only 21 genes were found to be downregulated in all of the tissue samples (Fig 1d). On comparing the genes associated with Fe homeostasis, it was observed that only 5 genes were commonly upregulated in roots and leaves of seedling and the mature stage. Similarly, 9 genes were expressed as downregulated genes (Fig 1e).

### 3.4 DEGs involved in ROS and Scavenging

ROS homeostasis greatly depends on the balance of ROS production and their scavenging. Under Fe^2+^ toxicity, stress-responsive, and source of H_2_O_2_ generation genes such as Respiratory burst oxidase homolog (Rboh) were differentially expressed in both roots and leaves (Fig 3a). Only *OsRbohH* was upregulated in roots, whereas *OsRbohF* in leaves of both growth stages. Simultaneously, enzymatic ROS scavenging enzymes such Superoxide dismutase (SOD), Catalase (CAT), Ascorbate peroxidase (APX), Guaiacol peroxidase (GPX), Glutathione reductase (GR), Peroxiredoxins (PRXs), Glutaredoxin (GRX) and Peroxidase (POX) were differentially expressed (Fig 3a). The expression of CAT was tissue and growth-specific. Only *OsCATA* was upregulated in roots, whereas *OsCatB* and *OsCATC* were upregulated in the leaves seedling and mature stage. Similarly, *OsSOD2* was upregulated in roots, whereas *OsSOD3* in leaves of both growth stages. Interestingly, *OsGR2* and *OsGR3* were commonly upregulated in all tissue conditions. Among various APX, *OsAPX1* and *OsAPX7* were upregulated in all tissue types. In roots, GPX were downregulated, whereas *OsGPX1*, *OsGPX2*, *OsGPX3* were upregulated in leaves of both stages. *OsPRXB* was highly upregulated in all tissue samples, where *OsPRXA* was specific in the leaves of seedlings. POX and non-enzymatic antioxidant Glutathione S-transferases (GST) are comprised of a large gene family and one of the highest DEGs in this study. Approximately 104 POX and 72 GST genes were differentially expressed across all tissue types. *OsPRX15* and *OsPRX93* were highly upregulated in roots whereas *OsPRX24* and *OsPRX30* in leaves seedling and *OsPRX113*, *OsPRX109*, *OsPRX110*, *OsPRX111*, etc. were highly upregulated in the mature stage. In roots, *OsGSTU44*, *OsGSTU43*, *OsGSTU47*, *OsGSTU8*, etc., were highly upregulated. Similarly, *OsGSTU30*, *OsGSTU19*, *OsGSTU36*, *OsGSTU12*, etc., were upregulated in leaves seedlings, whereas *OsGSTU8*, *OsGSTU6*, *OsGSTU17*, *OsGSTU24*, etc. were upregulated in the mature stage. These results suggested that the enzymatic pathways GST, POX gene families play a crucial role in protecting aromatic Keteki joha rice against oxidative damage caused due to Fe^2+^ toxicity.

**Figure 3:**
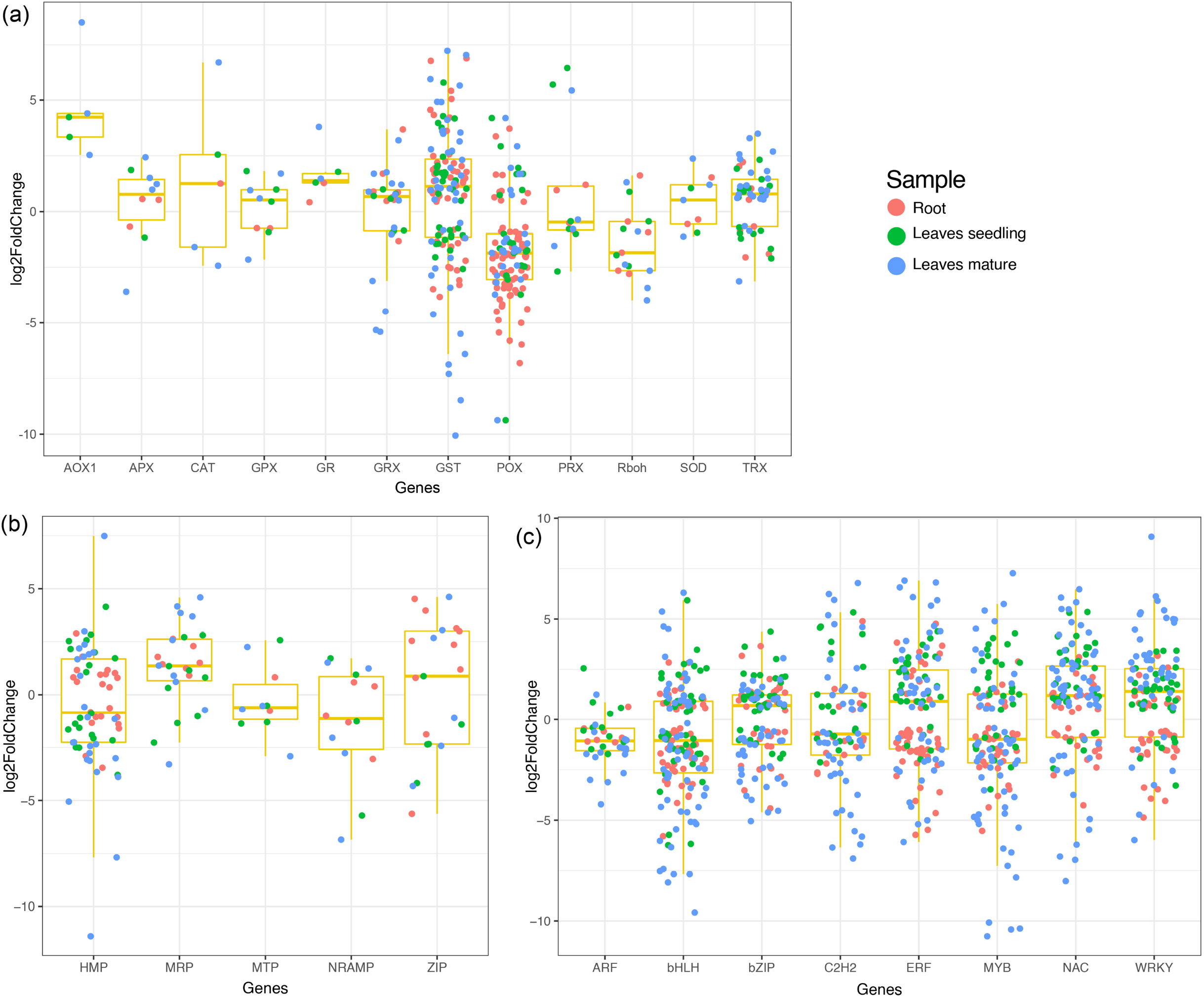
Boxplot showing the expression DEGs in different tissue samples. (a) Expression of antioxidant and ROS scavenging related DEGs; (b) Expression of heavy metal related DEGs and (c) Expression of transcription factors.

### 3.5 Differentially expressed transcription factors and heavy metal responsive genes under Fe^2+^ toxicity

Several transcription factors were differentially expressed under Fe^2+^ toxicity (Fig 3c). These include, Auxin response factors (ARF), Basic helix-loop-helix (bHLH), NAC (NAM, ATAF, and CUC), Myeloblastosis (MYB), Basic leucine zipper (bZIP), WRKY, APETALA2/Ethylene-responsive factor (AP2/ERF), C2H2 zinc-finger domain (C2H2) were prevalent (Fig 3c). Briefly, 100 bHLH, 70 bZIP, 121 NAC, 56 C2H2, 75 MYB, 81 AP2/ERF, and 69 WRKY transcription factors were differentially expressed in roots and leaves. Similarly, several metal transporters were differentially expressed under Fe^2+^ toxicity (Fig 3b). About 34 heavy metal-associated domains containing Heavy Metal-Associated Protein (HMP), 12 Multidrug resistance-associated protein (MRP), 7 Metal tolerance protein (MTP), 7 Natural resistance-associated macrophage protein (NRAMP), and 11 Zrt and Irt-like protein (ZIP) were differentially expressed.

### 3.6 DEGs related to abiotic stress and nutrient deficiency

Among the several DEGs, some well-known abiotic stress and nutrient deficiency-related genes were also observed. Auxin efflux carrier component genes were highly upregulated in roots where it was either downregulated or unexpressed in leaves (Fig S3). Abiotic stress-related genes 9-cis-epoxycarotenoid dioxygenase (*OsNCED4*/*OsNCED3*), Receptor-like Cytoplasmic Kinase 253 (*OsRLCK253*), phosphate dikinase (*OsPPDKA*), Cyclin-like F-box domain-containing protein (*OsMsr9*) were differentially expressed. In addition, Submergence Tolerance genes Pyruvate decarboxylase (*OsPDC4*), Phosphoenolpyruvate carboxykinase (*OsPEPCK*), Acireductone dioxygenase 1/ submergence-induced protein 2 (*OsARD1/OsSIP2*) and B12D protein were differentially expressed. Low nutrient responses genes *OsHB1*, *OsHB2* and *OsLPR2* were mostly upregulated (Fig S3).

### 3.7 Gene ontology and KEGG pathway affected by Fe^2+^ toxicity

Gene ontology (GO) is useful for describing the functions of gene products. GO terms of the DEGs in roots, leaves of seedling and mature stage were very similar. From comparative GO term analysis, top and common biological process GO terms include response to stress (GO:0006950), response to stimulus (GO:0050896), secondary metabolic process (GO:0019748), response to abiotic stimulus (GO:0009628), lipid metabolic process (GO:0006629), etc. (Fig 4). Similarly, the top and common molecular process GO terms include, catalytic activity (GO:0003824), drug binding (GO:0008144), oxygen binding (GO:0019825), transferase activity (GO:0016740), etc. (Fig S4). Besides top and common cellular components, GO terms of DEGs also belonged to cell (GO:0005623), external encapsulating structure (GO:0030312), cell wall (GO:0005618), extracellular region (GO:0005576), etc. (Fig S5).

**Figure 4:**
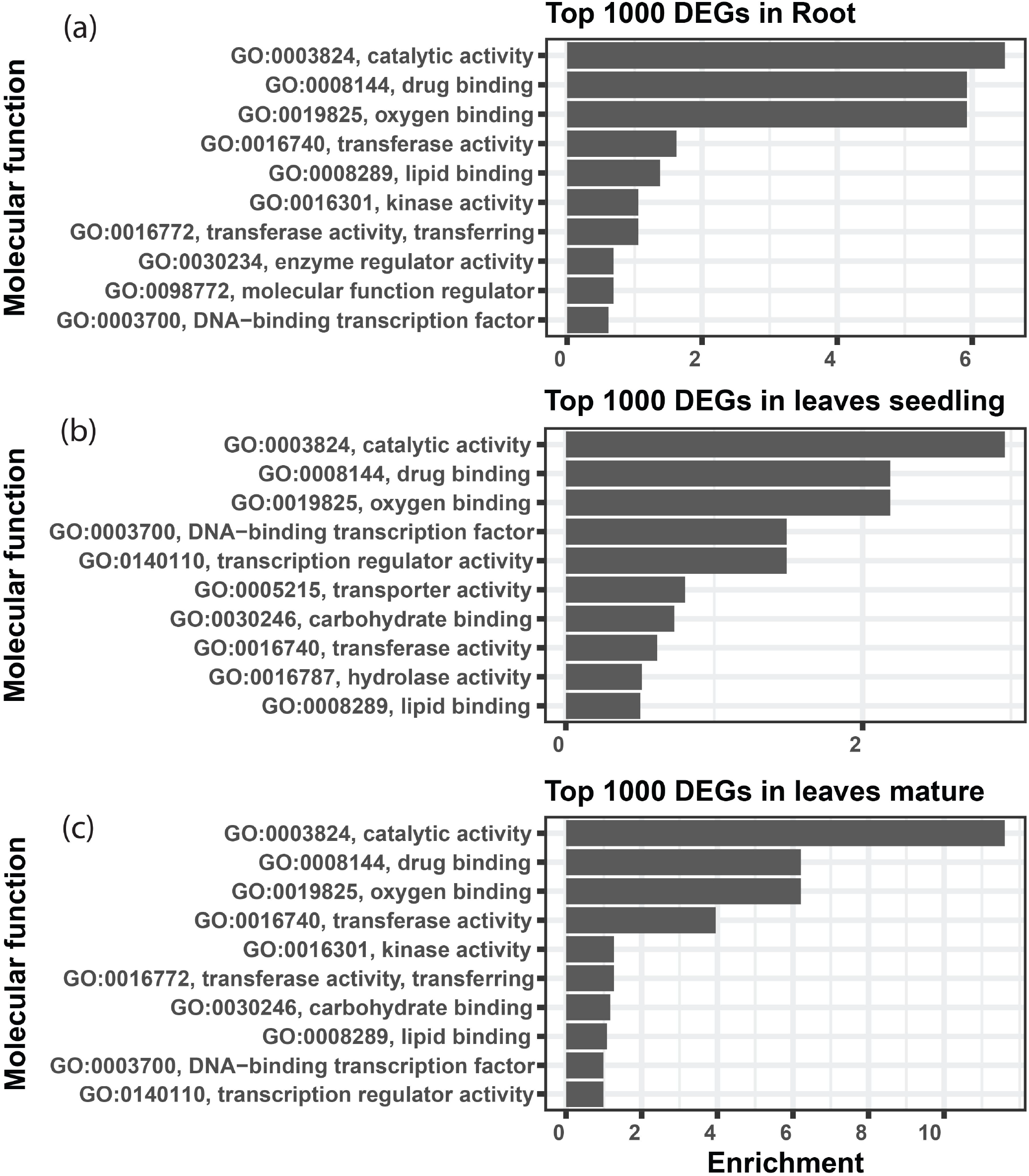
Gene ontology showing the biological process terms associated with top 1000 DEGs. The top 10 biological process terms were represented by applying Fishers exact statistical test and classic topGO package algorithm in R program. The ggplot2 package of R program was used to create the graphical representation.

The KEGG pathways of the DEGs were evaluated in the context of adaptation and responses to Fe^2+^ toxicity. Among the several pathways, significantly and differentially expressed pathways were evaluated by the pathview-web tool. Log2FoldChange value of each DEGs was integrated into their respective KEGG gene ID, and their active or passive state was evaluated and accordingly, multiple states pathways were presented (Fig 5). Top upregulated pathways were Biosynthesis of amino acids (osa01230), RNA degradation (osa03018), Spliceosome (osa03040), Glycolysis/ Gluconeogenesis (osa00010), Carbon metabolism (osa01200), Glutathione metabolism (osa00480) etc (Fig 5 and Fig 6). The most downregulated pathway includes Phenylpropanoid biosynthesis (osa00940), Photosynthesis (00195), Fatty acid elongation (osa00062), Ribosome (osa03010), DNA replication (osa03030), etc. (Fig 5 and Fig S6).

**Figure 5:**
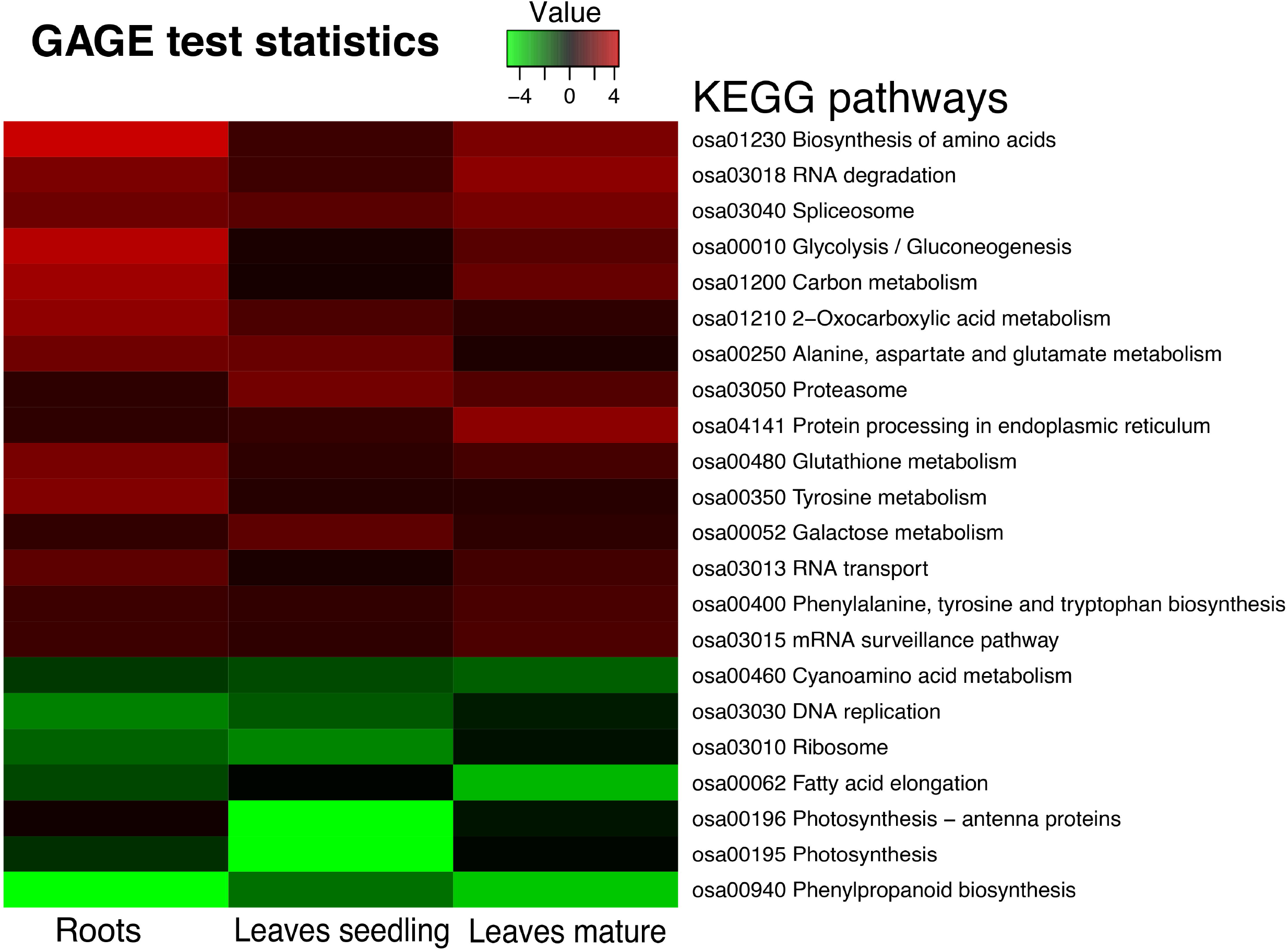
Heatmap showing the Kyoto Encyclopedia of Genes and Genomes pathways of the differentially expressed genes. GAGE statistics were applied, and the top significant pathways were represented using the pathview-web tool.

**Figure 6:**
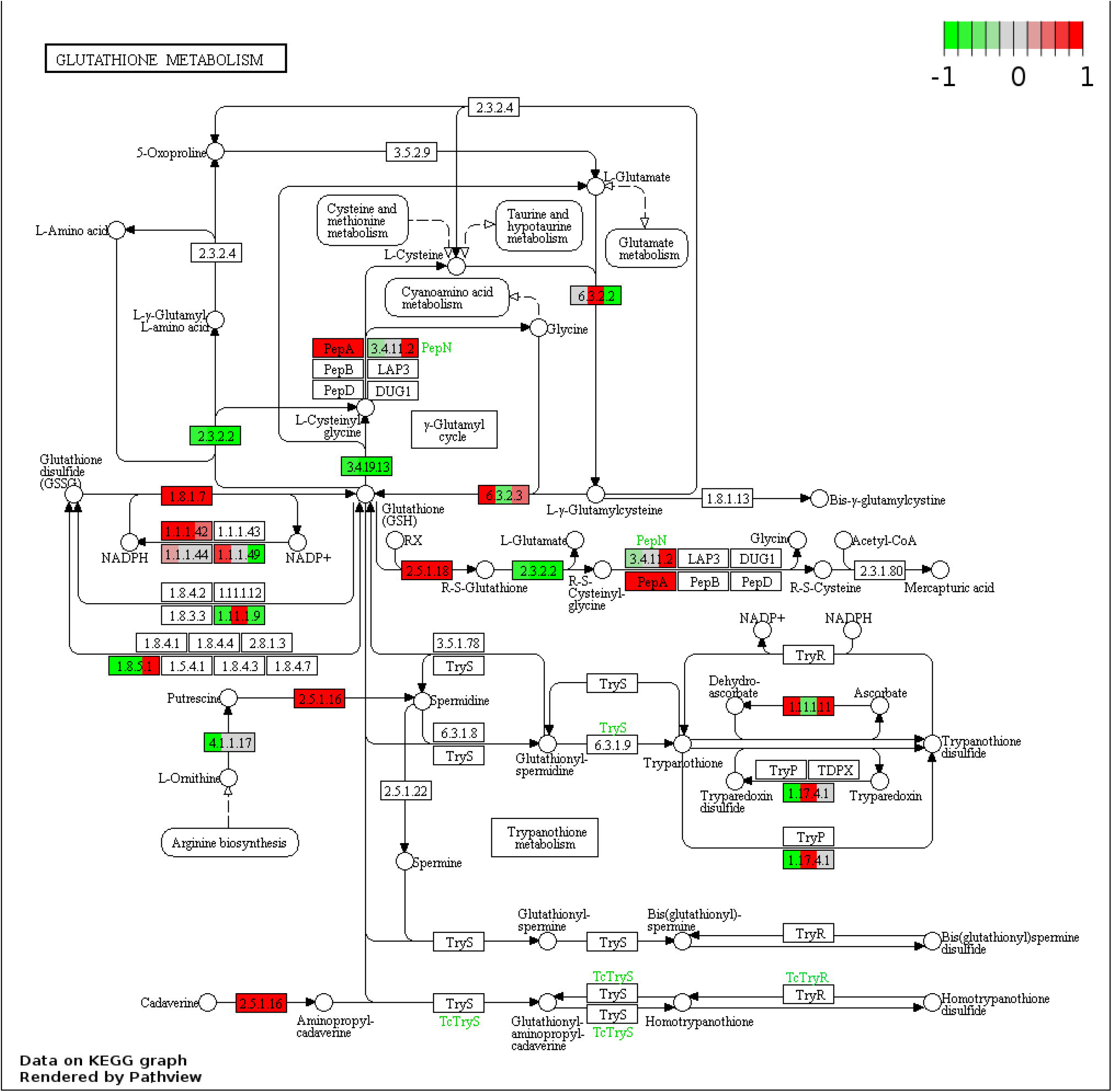
Representative figure showing the multiple states glutathione metabolism pathway induced under Fe^2+^ toxicity. DEGs involved in the pathway are highlighted in red (Upregulated) and green color (down-regulated).

### 3.8 Differential exon usage of genes

Differential exon usage of the DEGs was inferred in response to Fe^2+^ toxicity. About 173 in root seedling, 692 in leaves seedling and 1415 genes in leaves of the mature stage were identified to use different exons (Fig S7). Among them, Fe homeostasis genes were also identified to use different exons. In roots, *OsFRDL1*, *OsNRAMP1*, *OsVMT*, and *OsTOM3* used different exons between the control and treated groups. Similarly, the *OsFRDL1*, *OsIRO2*, *OsMIR* were in leaves seedling, and *OsFER1*/*2*, *OsSAM2*, *OsPEZ2*, *OsSAM1*, *OsIDEF2*, *OsTOM3*, and *OsVMT* used different exons between the control and treated groups (Fig S8).

### 3.9 Validation of gene expression

The expression of RNA-Seq data was validated by qRT-PCR analysis of representative genes (Fig 6). Under Fe^2+^ toxicity, expression of *Os*FER1/2 was upregulated in both root and leaves of the seedling stage. In roots, other induced genes include, *OsVMT*, *OsB12D*, and *OsGSTU44*. In contrast, similar to the RNA-Seq results, the *OsIRT1*, *OsYSL15*, *OsNAS1* and *OsTOM1* were downregulated. Similarly, the *OsYSL15* and *OsTOM1* were also downregulated in leaves of the seedling stage (Fig 7).

**Figure 7:**
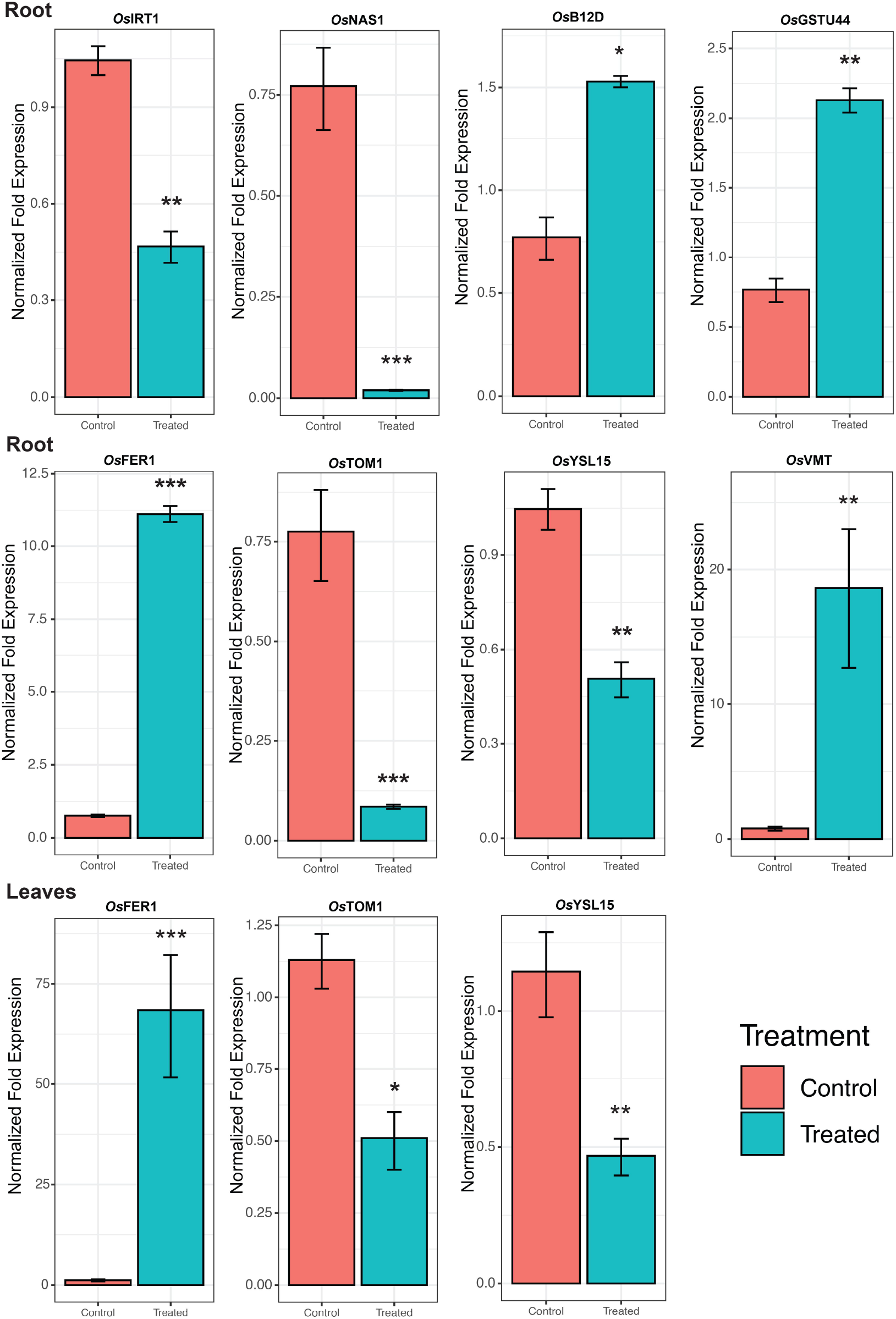
Expression of Fe homeostasis related genes. qRT-PCR confirmed RNA-seq results. Error bar shows the results of 3 biological replicates. The significant level indicates the significant difference in means between the control and treated group calculated by t-test. Significant codes with respect to P-values are *** 0.001, ** 0.01 and * 0.05.

A significant upregulation of ROS Scavenging genes and different abiotic stress-related TFs, genes serve to be the key mechanism involved against severe Fe^2+^ toxicity tolerance. Accordingly, a new defense mechanism is hypothesized that alleviates the excess Fe^2+^ in addition to the other mechanism of defense against Fe^2+^ toxicity (Fig 8 and Fig 9).

**Figure 8:**
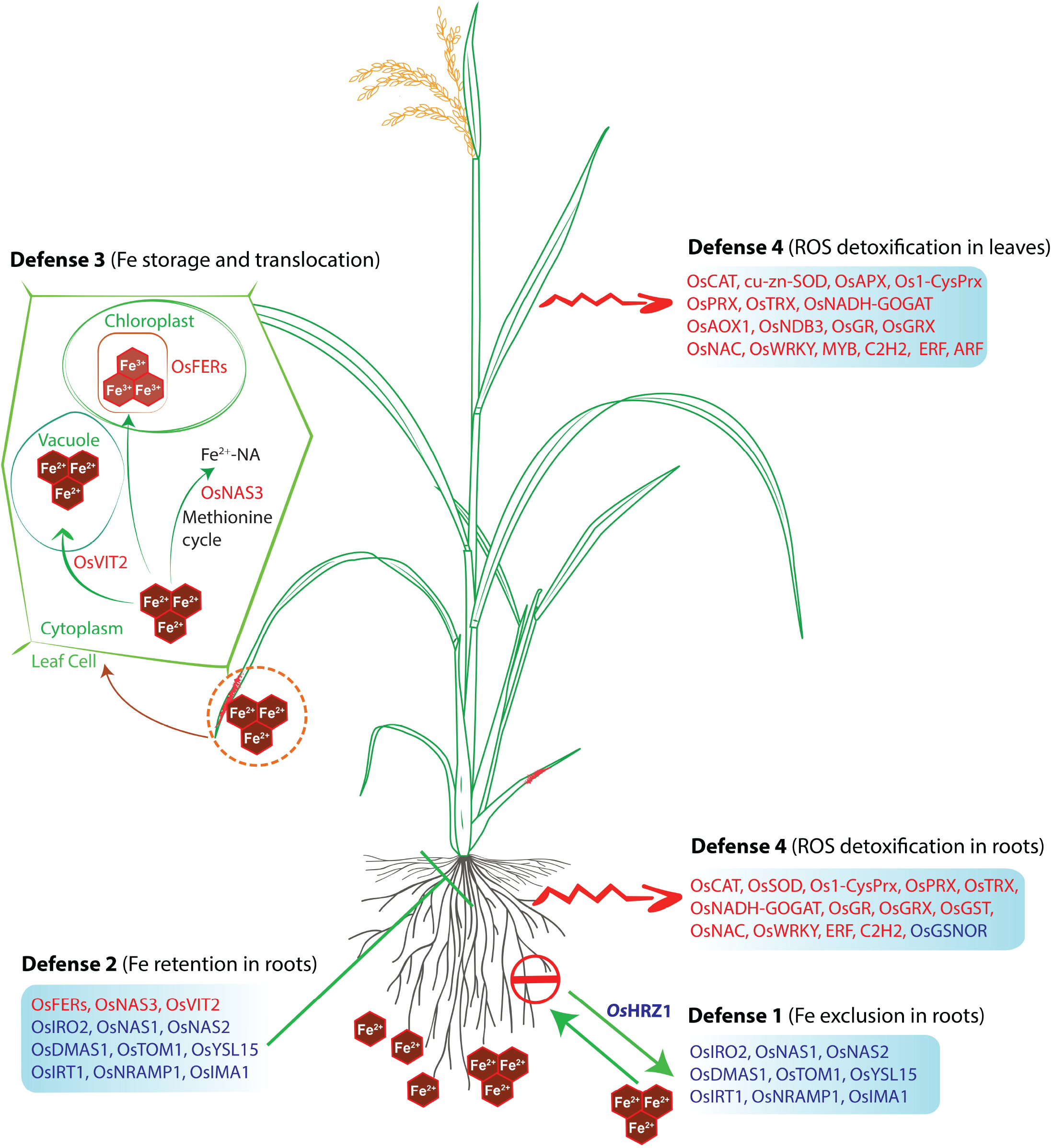
Hypothetical model of the defense mechanisms of rice against Fe toxicity. The model was adapted from Aung et al. (2020) and updated with our findings. Red letters indicate upregulated genes, whereas blue letters indicate downregulated genes under Fe-excess. The rice plant used in this model was adopted from the Gramene database (https://www.gramene.org/) and redrawn.

**Figure 9:**
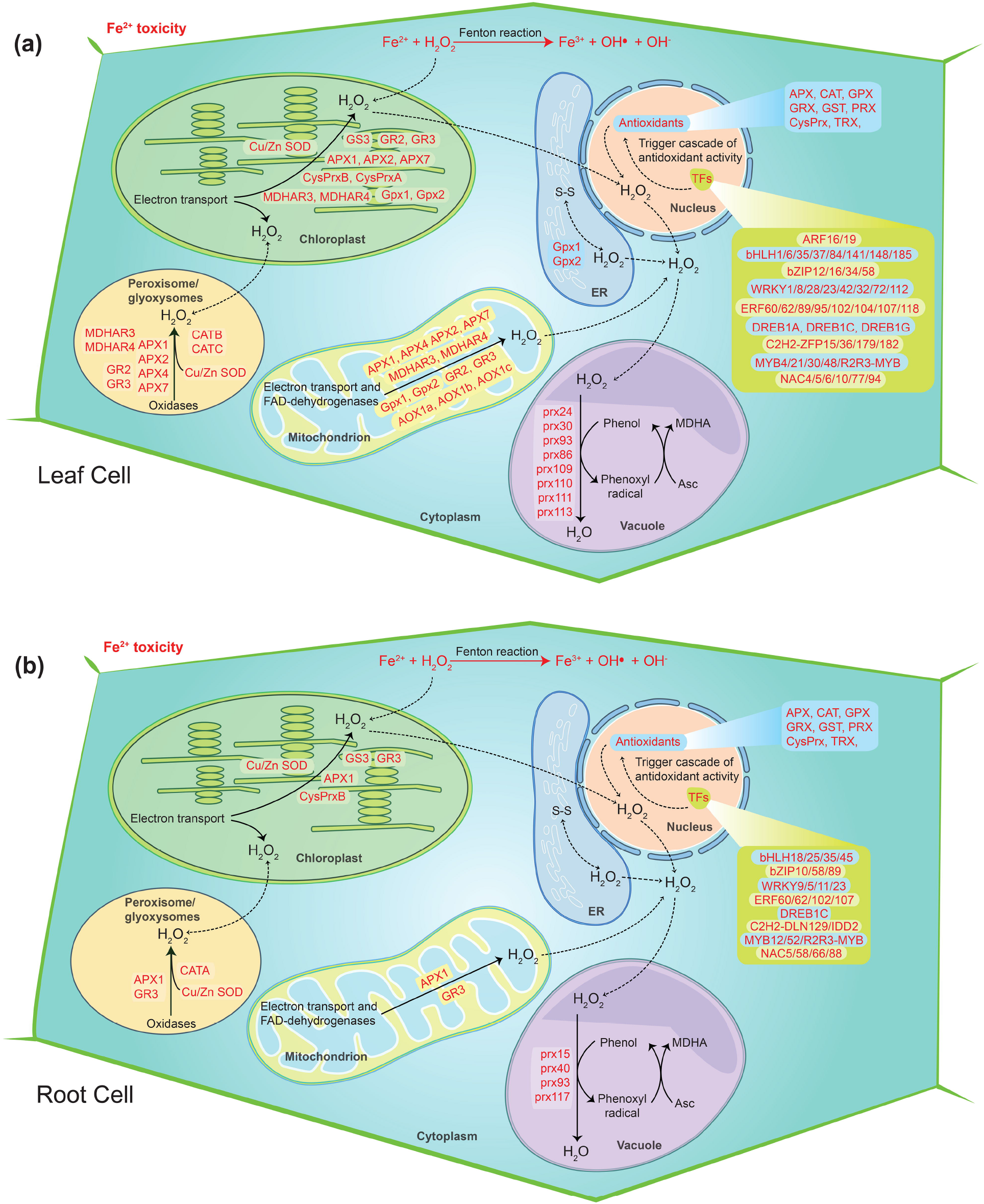
Hypothetical model of the cellular ROS detoxification mechanism under severe Fe^2+^ toxicity in rice.

## 4. Discussion

Fe^2+^ toxicity significantly inhibits the growth and productivity of rice. Fe^2+^ toxicity interrupts the metabolism and functioning of plants affecting root development and shoot inhibition ^11,13^. We have previously reported the effect of Fe^2+^ toxicity in aromatic Keteki Joha ^13^. In this study, we have performed a transcriptomic analysis of Keteki Joha under severe Fe^2+^ toxicity (2.5 mM) to investigate how it responds under stress conditions. Fe homeostasis genes were well characterized in Fe deficient condition ^35,36^; however, their role is still unclear in Fe excess. In this study, nearly 69 Fe homeostasis-related genes and their homologs were identified, which were mostly downregulated. Our RNA-Seq results share similarities with the previous microarray result with several other exceptions ^11,37^. Thus, these findings will help to further ameliorate the understanding he Fe homeostasis in rice. Differential exon usages of few Fe homeostasis genes were identified. From differential exon usage, possible alternative splicing of DEGs can be inferred ^31^. However, validation of the predicted genes will be essential to conclude any transcriptional alteration or changes under Fe^2+^ toxicity.

Four different types of tolerance mechanisms have been reported in rice ^12^. However, Fe sensing is crucial at the initial response stage in Fe homeostasis gene expression and regulation based on Fe availability ^38^. *OsIDEF1*, *OsIDEF2*, *OsIRO2* positively regulate Fe deficiency-inducible genes involved in DMA-based Fe(III) acquisition, Fe^2+^ uptake, and Fe translocation. Additionally, *Os*IDEF1 was directly bound with Fe^2+^ and other divalent metal ions, suggesting an intracellular Fe sensor ^39^. Moreover, *OsHRZ1* and *OsHRZ2* can also bind with Fe via hemerythrin domains, thus characterized as another Fe sensor candidate. Their knockouts have negatively modulated the response to Fe-deficiency in rice. In rice, *OsIDEF1* controls the expression of both *OsHRZ1* and *OsHRZ2* ^40^. However, in this study, the *OsIDEF1* gene was not found to be differentially expressed among all the tissues. *OsHRZ1* being downregulated, suppression of Fe transporter genes in transcriptional level are quite unlike under severe Fe^2+^ conditions. Besides, *OsHORZ1*, a hemerythrin domain-containing protein, was also downregulated, which is known to repress *OsHRZ* functions ^38,39,41^. In this context, further study of transcription factors in contrasting genotypes may provide a better understanding of their downstream regulation and proper functioning under Fe^2+^ toxicity. However, Fe-excess responses in rice plants are thought to be partially independent of Fe-deficiency ^38^. A positive Fe-deficiency regulator *OsPRI1*, a target gene of *OsHRZ1* was also unexpressed in this study. Despite that, *OsNAS3* was upregulated, which is important for Fe detoxification in the root ^42^. In addition, *OsFRDL1* was also upregulated, which is responsible for root-to-shoot Fe translocation and is believed to play a significant role in the distribution of Fe into old leaves. Thus, *OsFRDL1* is crucial for the minimization of Fe^2+^ toxicity in roots. As a result, *OsFER1/OsFER2* and *OsVIT2* were also highly upregulated in leaves, thus defending the Fe overload. Overall, our results are quite supported by the proposed Fe-excess defense mechanisms ^12^. In exception, *OsVMT* was highly upregulated in both root and leaves, and it is known to be expressed where Fe and Zn are highly deposited. Knockout of these genes enhances Fe and Zn accumulation in polished rice grains as DMA increases solubilization of Fe and Zn deposited in the node ^32^. Considering the function and expression of *OsVMT* under Fe-excess conditions, it is likely to involve in Fe detoxification. Interestingly, the *OsTOM3* DMA-efflux transporter gene belonging to Zinc induced facilitator (ZIFL subfamily) was upregulated in all the tissues, thereby involved in metal transport. The characterization of this gene needs to be further studied for better clarification about its role in Fe^2+^ toxicity. The Phenolic Efflux Zero1 induced in root tissue plays a significant role in the efficient translocation of protocatechuic acid (PCA) and caffeic acid from roots to shoots in rice ^43^. Previously, it was found that expression of mitochondrial iron transporter (*OsMIT*) increases under Fe excess which is in accordance with our results, essential for the proper growth and development of rice ^36^. As reported earlier, *OsYSL9* is involved in iron translocation, particularly from endosperm to embryo in developing seeds ^44^. Yellow Stripe 1-Like (YSL) family transporters are responsible for the transport of metal-phytosiderophores; however, their role is unclear in rice. *OsYSL9* localizes in the plasma membrane and is believed to participate in the transportation of both iron (II)-nicotianamine and iron (III)-deoxymugineic acid into the cell. The expression of *OsYSL10* is similar to *OsYSL9* and may also participate in a similar mechanism. *OsYSL13* was expressed in roots, and mature leaves participate in the distribution of Fe from older leaves to new leaves ^45^. Earlier it was found that *OsNRAMP2* was highly induced in shoots and participate in Cadmium accumulation ^46^. Various studies have already been identified the role of *OsNRAMP2* to participate in Fe accumulation which is found to be upregulated in all the tissues. Acireductone dioxygenase, *OsARD1* is a metal-binding enzyme and is involved in the production of methionine as it binds with Fe^2+^ and catalyzes the formation of 2-keto-4-methylthiobutyrate (KMTB). In our study, *OsARD1* showed high upregulation in all the tissue types that might enhance rice tolerance under Fe^2+^ toxicity. Overexpression of *OsARD1* also leads to the tolerance of abiotic stresses like submergence, drought and salinity ^47^.

ROS scavenging remains a key mechanism under severe Fe^2+^ toxicity among the different Fe-excess defense mechanisms, which needs a better understanding ^12^. Recently, few ROS scavenging genes, transcriptome factors and transporter genes were highlighted as probable candidates for the ROS scavenging system ^10–12,37^. However, specific ROS scavenging genes and their molecular mechanism are still unknown. Based on RNA-Seq transcriptome, we explored the possible ROS scavenging system in rice under severe Fe^2+^ toxicity. Accordingly, a new defense mechanism (defense 4), i.e., ROS detoxification has been hypothesized as a possible mechanism of tolerance against severe Fe^2+^ toxicity. Excess Fe^2+^ is very lethal to plants under acidic conditions as they catalyze H_2_O_2_ and produce a highly toxic hydroxyl free radical, term as Fenton reaction ^48^. Toxic hydroxyl free radical causes lipid peroxidation and promotes programmed cell death ^49^. Lipid peroxidation and programmed cell death can occur under different environmental stresses. However, Fe-mediated lipid peroxidation and programmed cell death results are quite different from others and are termed Ferroptosis ^50^. Rice has developed several Fe tolerance mechanisms, but at severe conditions, only the ROS homeostasis was hypothesized to be essential and important in the tolerance mechanism ^11,12^. Under Fe^2+^ toxicity, higher production of H_2_O_2_ was reported ^11,13^. H_2_O_2_ can be synthesized mostly in every cellular compartment, *viz*. peroxisomes, chloroplasts, mitochondria, nucleus, vacuoles, etc. CAT was reported to be mainly restricted only in peroxisomes with a very high concentration, thus removing H_2_O_2_ ^51^. In concordance with previous reports, all three CAT genes were upregulated in roots and leaves of seedling and mature stages. Further, the involvement of these particular CATs was confirmed by KEGG analysis.

Most of the APX, ferredoxin and thiol-based peroxidase such PRX genes were upregulated in both roots and leaves which are important for the removal of H_2_O_2_ in chloroplast using NADPH and photosynthetic electron transport via ferredoxin (2Fe-2S, iron-sulfur cluster binding domain-containing protein) as the ultimate reductant ^51^. Several POX genes were upregulated, which are known to be responsible for scavenging H_2_O_2_ by oxidizing various secondary metabolites. Removal of mitochondrial H_2_O_2_ requires a series of enzymatic detoxification processes. Mn-SOD scavenges superoxide radicals formed via mitochondrial electron transport chain (ETC). However, *Mn-OsSOD1* was not found to be differentially expressed in roots, whereas it was fractionally upregulated in leaves of both stages. The remaining H_2_O_2_ is detoxified by the PRX-TRX system or by the enzymes of the ascorbate-glutathione cycle. *Os1-CysPrxB* is highly expressed across all tissue types. Several TRX genes were also upregulated such as *OsTRX24*, *OsTRX23*, *OsTRX2*, etc.; thus, PRX-TRX probably regulates the detoxification of H_2_O_2_ in mitochondria under Fe^2+^ toxicity. GSTs were one of the most differentially expressed genes, and KEGG analysis revealed the glutathione pathway as one of the most actively regulated in this study. Glutathione synthase *viz. OsNADH-GOGAT1* and *OsNADH-GOGAT2* were identified as highly upregulated genes in roots involved in the glutathione synthesis pathway, whereas *OsNADH-GOGAT1* was identified in leaves. In addition, alternative respiratory pathway component genes like alternative oxidase (AOX) and NAD(P)H dehydrogenase (NDs) plays an essential role in minimizing ROS production ^52–55^. *OsAOX1a*, *OsAOX1b*, *OsAOX1c*, and *OsNDB3* were highly upregulated in leaves, thus suggesting the existence of the mechanism under Fe^2+^ toxicity. As described previously, the modulation of this mechanism was observed under cold stress in rice ^56^. Overall, expression of GSH, APX, MDHAR, DHAR, and GR indicates the active role of the ascorbate-glutathione cycle under Fe^2+^ detoxification which has been also affirmed by the KEGG pathway analysis.

Iron homeostasis is a complex network that is regulated by various transcription factors. The tale of transcription factors involved in the regulation of different Fe-acquisition was well characterized in *Arabidopsis*, whereas it is comparatively less explored in rice. In *Arabidopsis*, 16 bHLH transcription factors were described to be involved in Fe deficiency responses ^57^. In this study, several other bHLH TFs were differentially expressed, which need to be further elucidated. Moreover, in *Arabidopsis*, MYB, WRKY, AP2/ERF, and C2H2 TFs were involved in Fe acquisition, translocation, inhibition, and modulation of other Fe homeostasis genes ^57^. In rice, *OsIDEF1* (ABI3/VP1 TF family) and *OsIDEF2* (NAC TF family) *OsARF16* (ARF family) reported to modulate the expression of Fe-related genes and integrate auxin signals respectively, thus play a critical Fe homeostasis in rice ^57–59^. Apart from the above TFs, several other TFs were differentially expressed under Fe^2+^ toxicity, which might involve Fe homeostasis regulation in rice. Furthermore, TFs are well known to regulate various abiotic stress tolerance in plants. Recently, NAC (*OsNAC4*, *OsNAC5*, and *OsNAC6*) and WRKYs TFs were hypothesized to regulate severe Fe^2+^ tolerance in rice ^12^. Besides, S-nitrosoglutathione-reductase (GSNOR) fractionally downregulated in this study was reported to promote root tolerance to Fe toxicity *via* a nitric oxide pathway ^60^. WRKY transcription factors are well known to play a key role in both biotic and abiotic stress tolerance as well as plant hormones signal transduction and the MAPK signaling cascade in plants ^61–64^ Among the several upregulated WRKY TFs, *OsWRKY8* and *OsWRKY71* were involved in various biotic and abiotic stresses in rice ^63,64^. Moreover, APETALA2 /Ethylene Response Factors (AP2/ERF) are well known to regulate numerous abiotic stresses ^65,66^. Several AP2/ERF transcription factors were upregulated under Fe^2+^ toxicity. Interestingly, dehydration-responsive element-binding proteins (DREBs) transcription factors were downregulated in roots, whereas upregulated in leaves. Overall, differentially expressed TFs under Fe^2+^ toxicity is very intense, which need to be further studied to elucidate their specific role in understanding Fe^2+^ toxicity in rice. However, identification of their downstream target genes will remain important to uncover the possible signaling pathways in rice.

Comparative KEGG analysis revealed that genes involved in the Mitogen-activated protein kinases (MAPK) signaling pathway are highly upregulated in leaves under Fe^2+^ toxicity. MAPK signaling pathway is a common defense response of plants for various abiotic stresses ^67,68^. Along with several MPK genes, *OsMPK3* was upregulated in leaves which is known as a stress tolerance gene ^69^. In addition, pathogen-related protein 1 (*OsPRI1*) was also upregulated in all tissue types which was reported to play important roles in the plant metabolism in response to biotic and abiotic stresses ^70^.

## Conclusion

In this study, based on RNA-Seq, we identified the possible candidate genes that help in better understanding of Fe excess adaptation in rice. ROS scavenging was found to be the key mechanism involved in the detoxification of severe Fe^2+^ toxicity. As per our findings, upregulation of numerous ROS scavenging genes and various abiotic stress-related TFs under severe Fe^2+^ toxicity suggested that rice plants use the defense 4 mechanism known as ROS detoxification. Thereby, a new hypothetical model has been reconstructed in addition to the defense mechanism reported earlier against Fe^2+^ toxicity. Thus, our findings contribute to a better understanding of Fe homeostasis related genes and Fe-mediated ROS detoxification in rice.

## Additional information

The author(s) declare no competing financial interests.

## Acknowledgment

The author highly acknowledges Ministry of Tribal Affairs, Government of India for financial support under National Fellowship Scheme for Higher Education of ST Students, 2016-17. Award no: F1-17.1/2016-17/NFST-2015-17-STASS-2810

## Supplementary Figures

Figure S1: Representative photograph showing the growth of Keteki Joha under control and Fe Excess (2.5 mM) conditions.

Figure S2: Sample details for RNA Sequencing. A minimum of 3 biological replicates was used for sequencing.

Figure S3: Heatmap of known abiotic stress-responsive DEGs. ComplexHeatmap package of R program was used to prepare the heatmap. The color scale bar represents the log2FoldChange value of the DEGs.

Figure S4: Gene ontology molecular function terms associated with top 1000 differentially expressed genes. The top 10 biological process terms were represented by applying Fishers exact statistical test and classic topGO package algorithm in R program. The ggplot2 package of R program was used to create the graphical representation.

Figure S5: Gene ontology cellular component terms associated with top 1000 differentially expressed genes. The top 10 biological process terms were represented by applying Fishers exact statistical test and classic topGO package algorithm in R program. The ggplot2 package of R program was used to create the graphical representation.

Figure S6: Representative figures showing the differential expression of Glycolysis / Gluconeogenesis, MAPK signaling pathway and Photosynthesis KEGG pathways under severe Fe^2+^ toxicity. Multiple states pathway was rendered by Pathview-web tool and the DEGs involved in the pathway are highlighted in red (Upregulated) and green color (down-regulated).

Figure S7: Venn diagram showing the number of common differential exon usage DEGs in different tissues under Fe^2+^ toxicity.

Figure S8: Representative figure showing the differential exon usage of *OsFRDL1*. Significant differential exons usages are highlighted.

## Supplementary Tables

Table S1: Primers for qRT-PCR.

Table S2: RNA-Seq samples showing the number of reads and percentage of uniquely mapped reads into the rice genome.

