## Supplementary Figures for "Transcriptomic analysis revealed reactive oxygen species scavenging mechanisms associated with ferrous iron toxicity in aromatic Keteki Joha rice"

Preetom Regon^1^

<https://orcid.org/0000-0002-6862-5379>

Sangita Dey^1^

<https://orcid.org/0000-0001-6680-1167>

Mehzabin Rehman^2^

<https://orcid.org/0000-0002-0419-360X>

Amit Kumar Pradhan^2^

<https://orcid.org/0000-0003-1705-3347>

Bhaben Tanti^2^

<https://orcid.org/0000-0002-7594-4562>

Anupam Das Talukdar

<https://orcid.org/0000-0001-8916-2791>

Sanjib Kumar Panda^3^

<https://orcid.org/0000-0001-5131-9693>

**Author affiliation**

^1^Department of Life Science and Bioinformatics, Assam University, Silchar-788011, Assam, India.

^2^Department of Botany, Gauhati University, Guwahati-781014, Assam, India.

^3^Department of Biochemistry, Central University of Rajasthan, Ajmer- 305817, India.

Corresponding

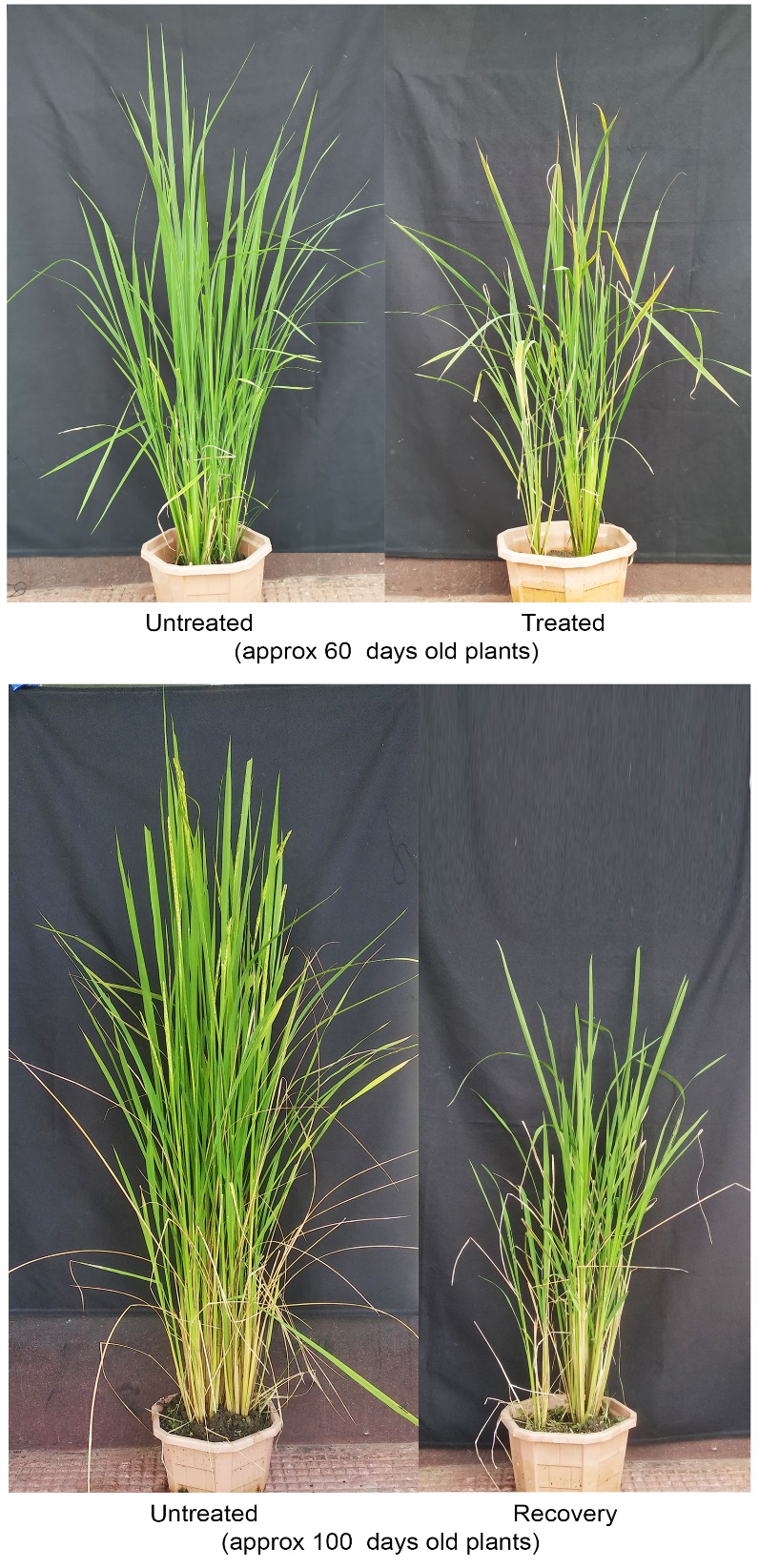

Fig S1: Representative figure showing the morphology of Keteki Joha grown under control and Fe Excess (2.5 mM) conditions.

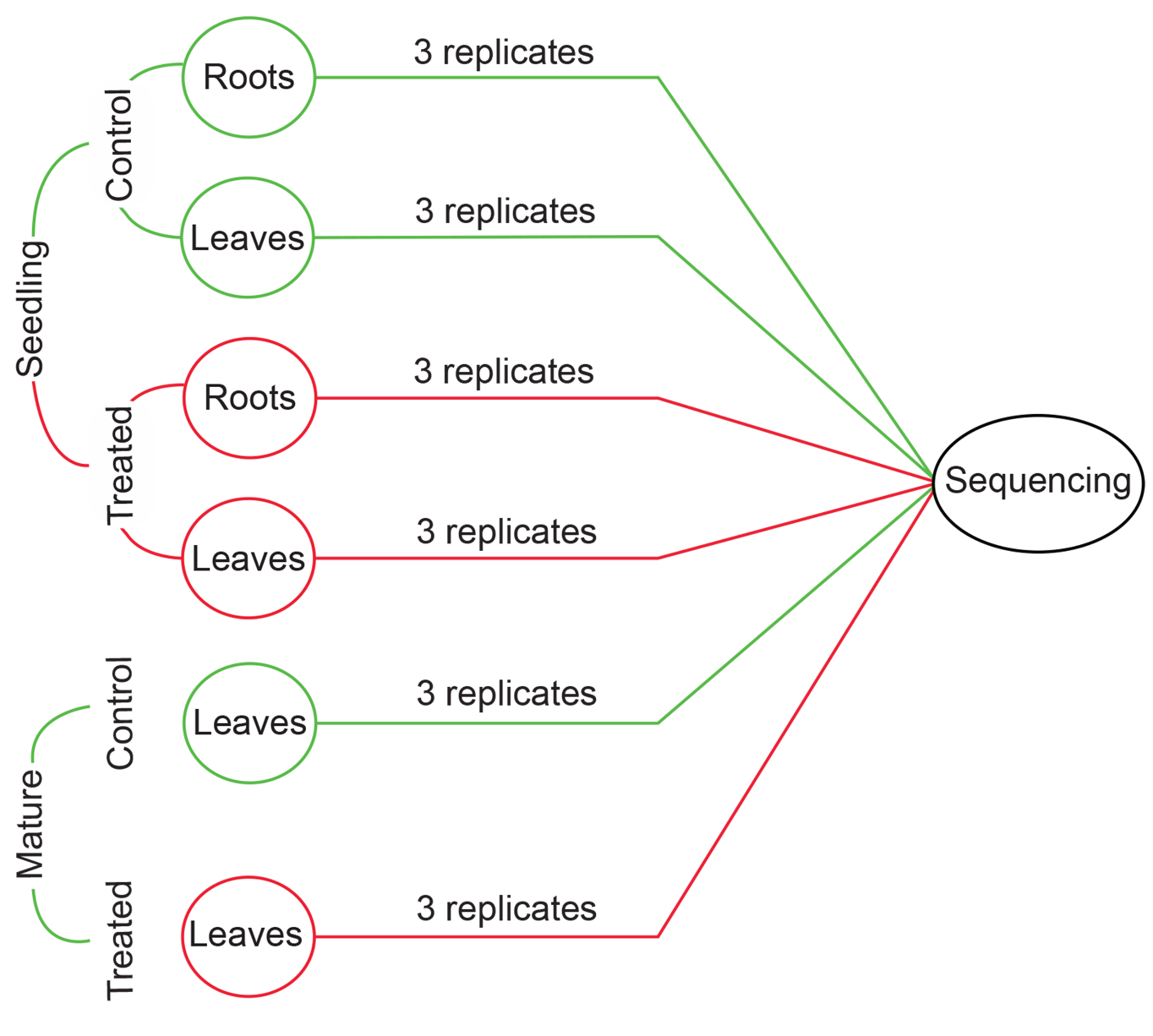

Figure S2: Sample details for RNA Sequencing. A minimum of 3 biological replicates was used for sequencing.

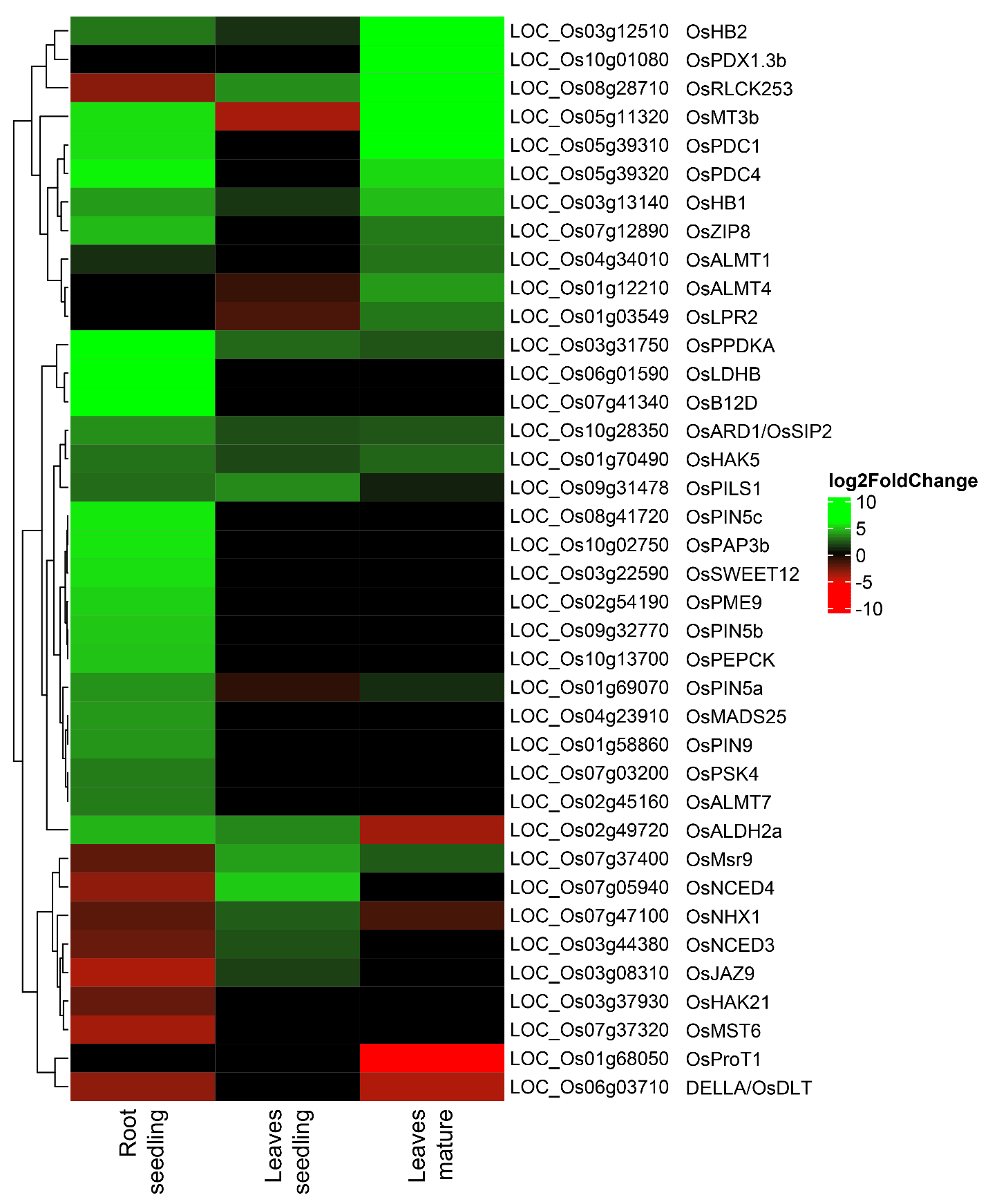

Figure S3: Heatmap of known abiotic stress-responsive DEGs. ComplexHeatmap package of R program was used to prepare the heatmap. The color scale bar represents the log2FoldChange value of the DEGs.

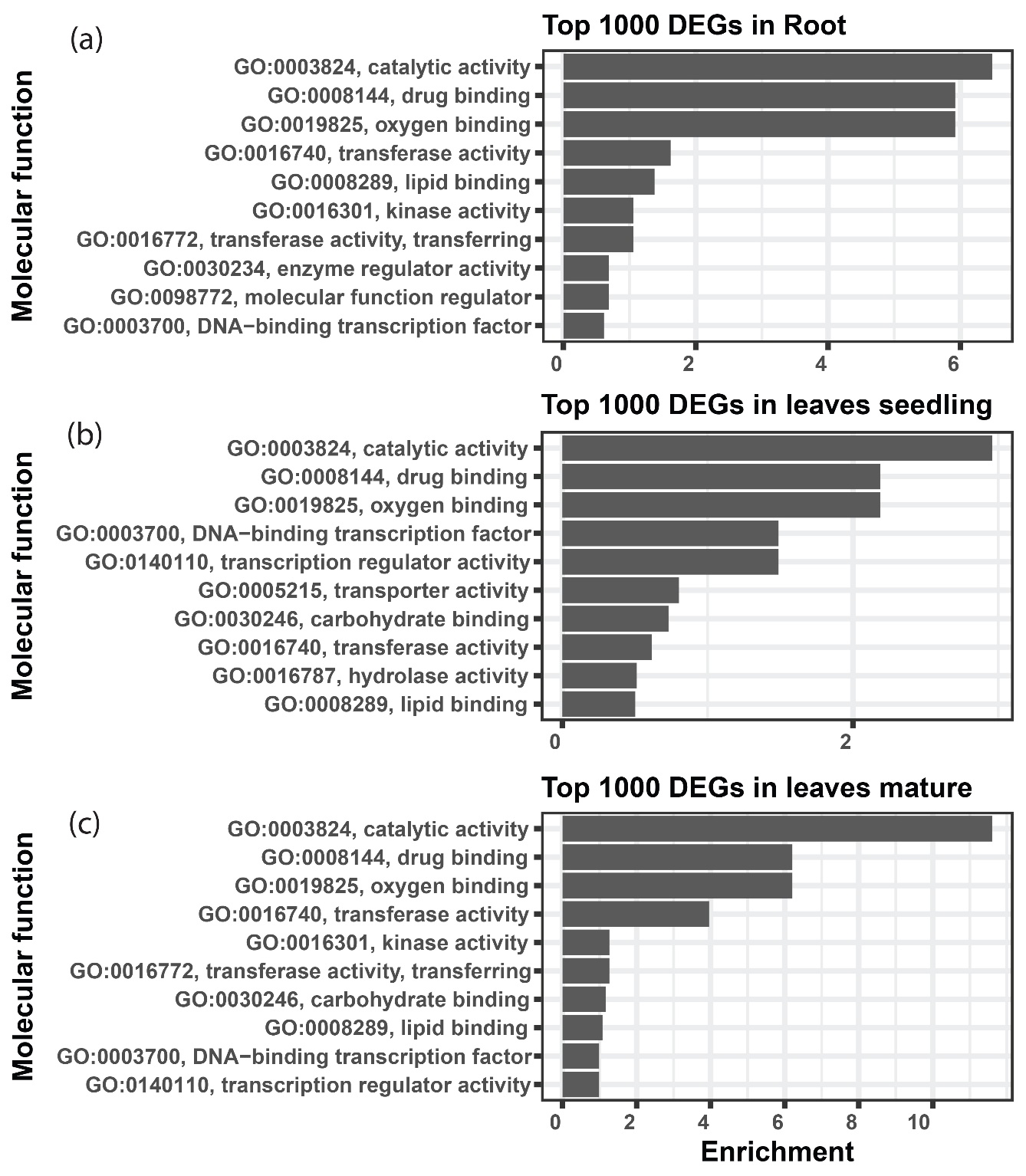

Figure S4: Gene ontology showing the Molecular function terms associated with top 1000 DEGs. The top 10 biological process terms were represented by applying Fishers exact statistical test and classic topGO package algorithm in R program. The ggplot2 package of R program was used to create the graphical representation.

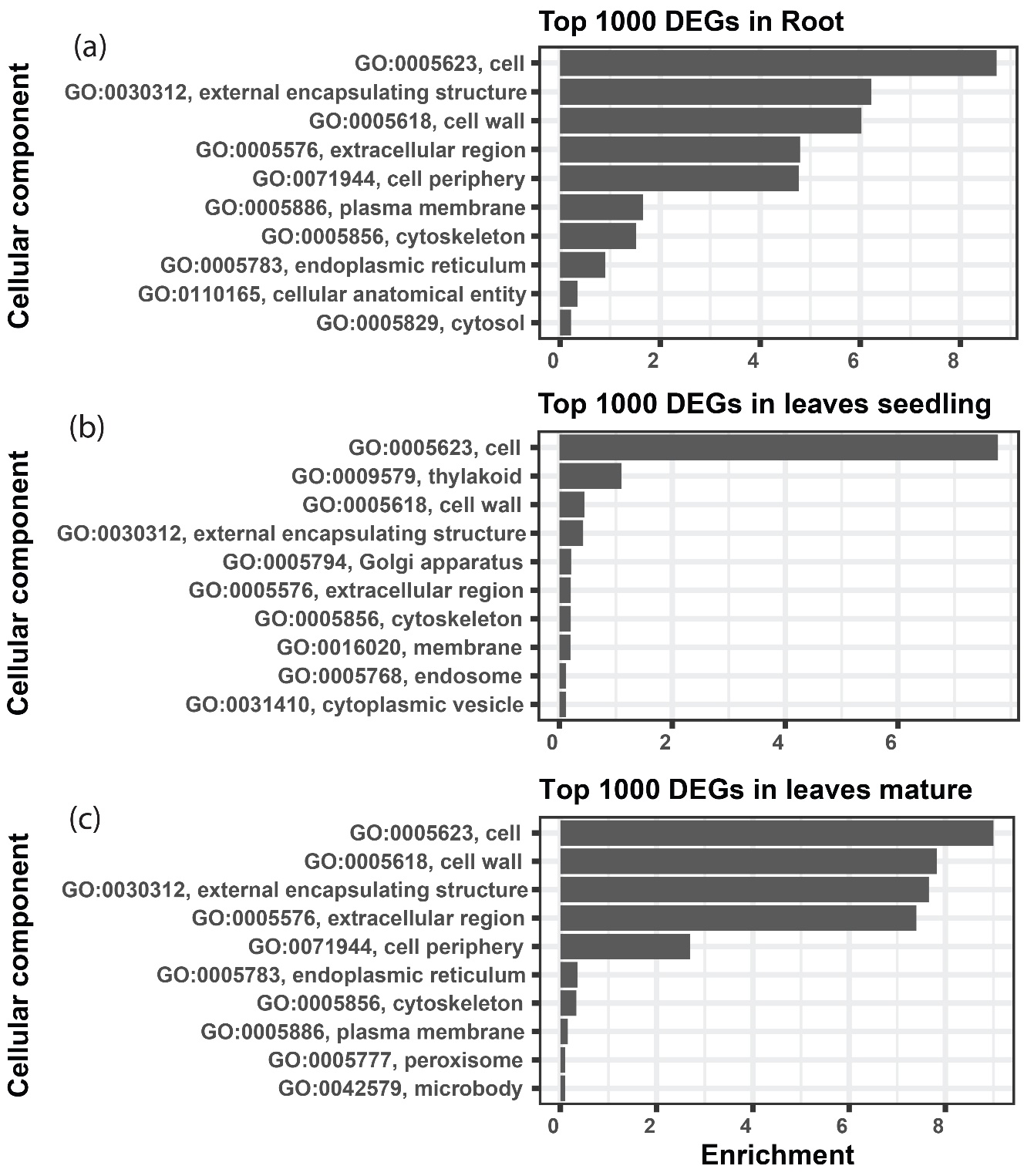

Figure S5: Gene ontology showing the Cellular component terms associated with top 1000 DEGs. The top 10 biological process terms were represented by applying Fishers exact statistical test and classic topGO package algorithm in R program. The ggplot2 package of R program was used to create the graphical representation.

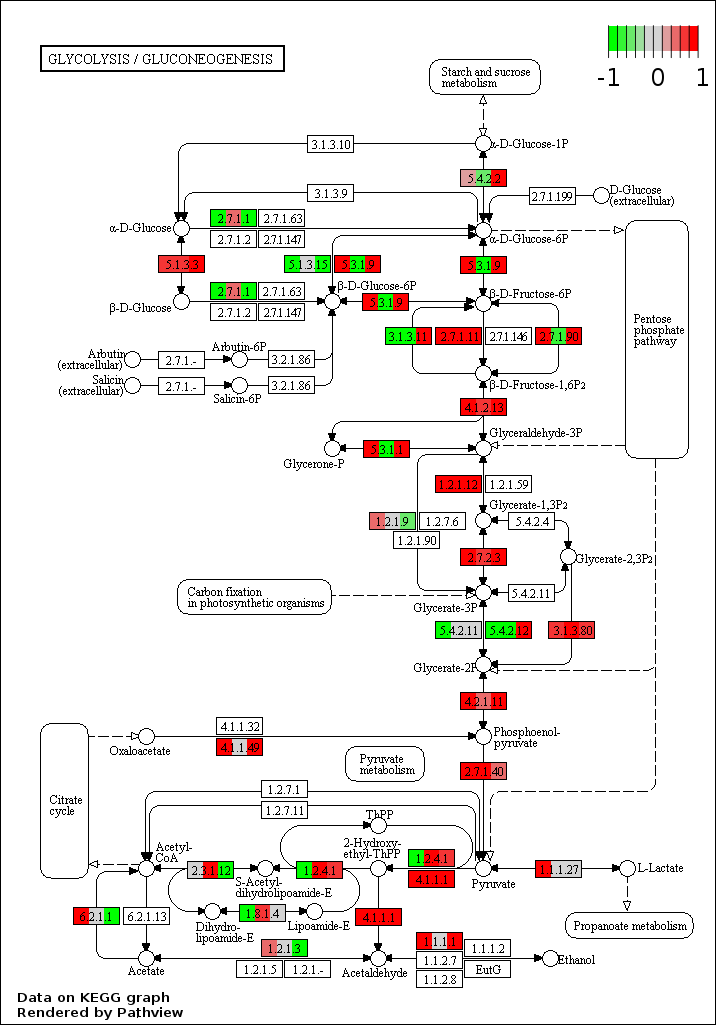

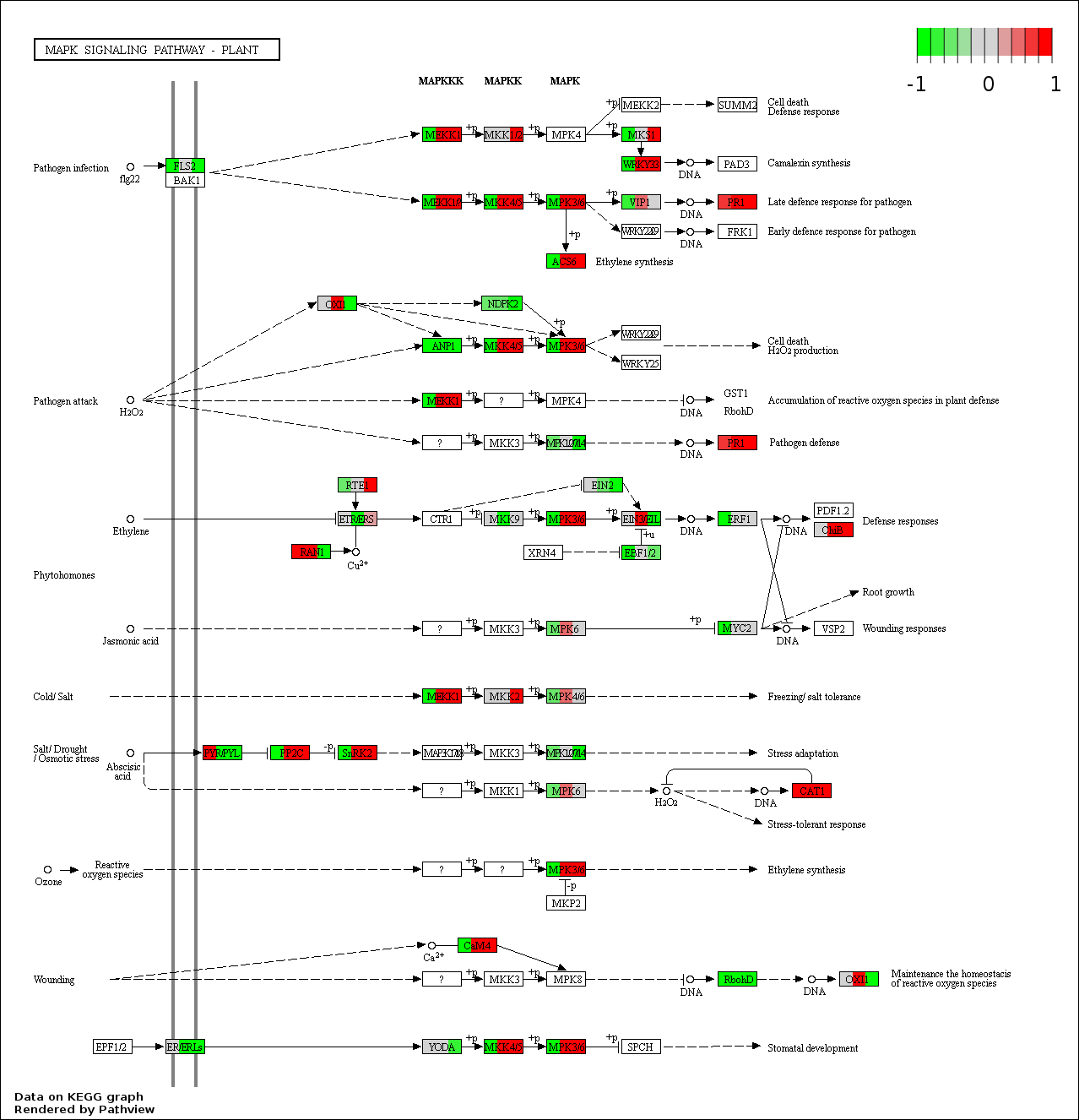

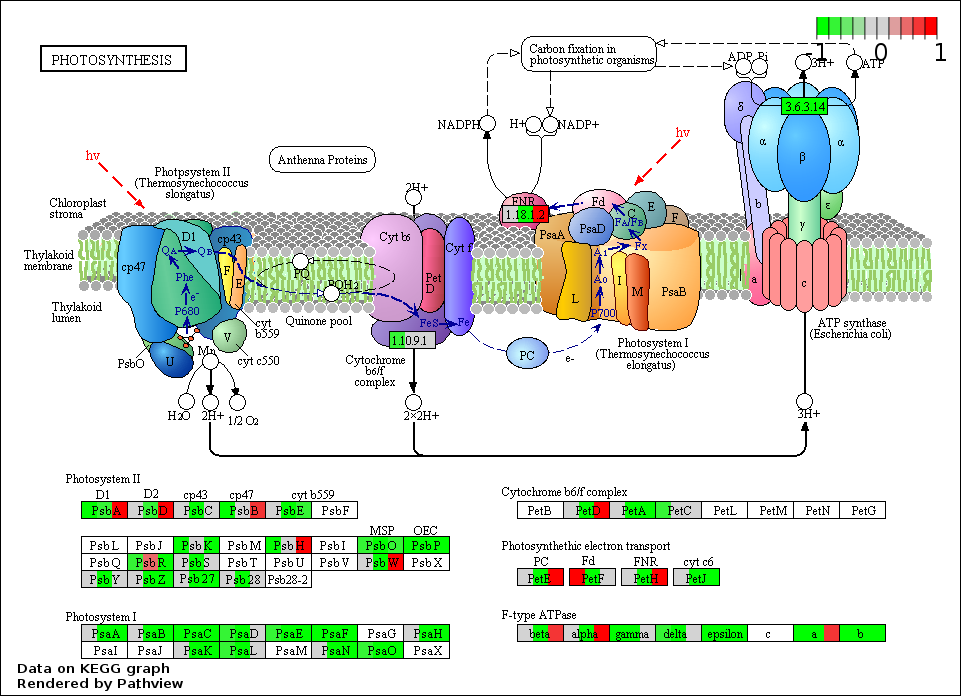

Figure S6: Representative figures showing the differential expression of Glycolysis / Gluconeogenesis, MAPK signaling pathway and Photosynthesis KEGG pathways under severe Fe^2+^ toxicity.

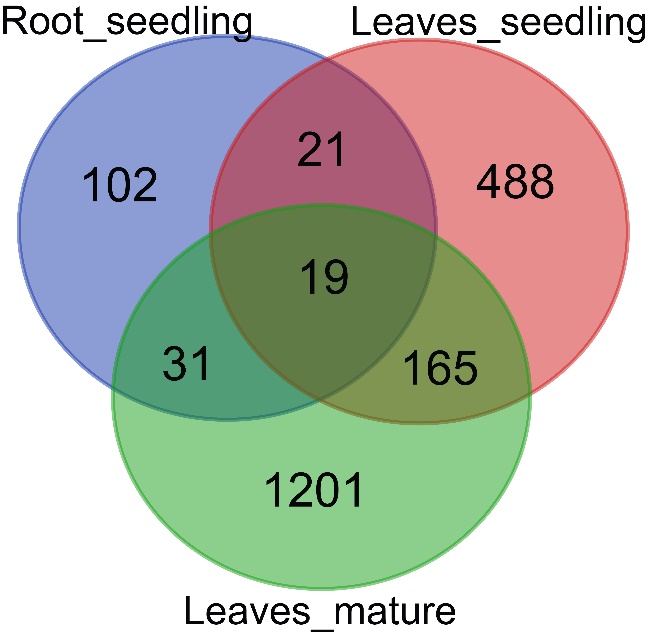

Figure S7: Venn diagram showing the number of common differential exon usage DEGs in different tissues under Fe^2+^ toxicity.

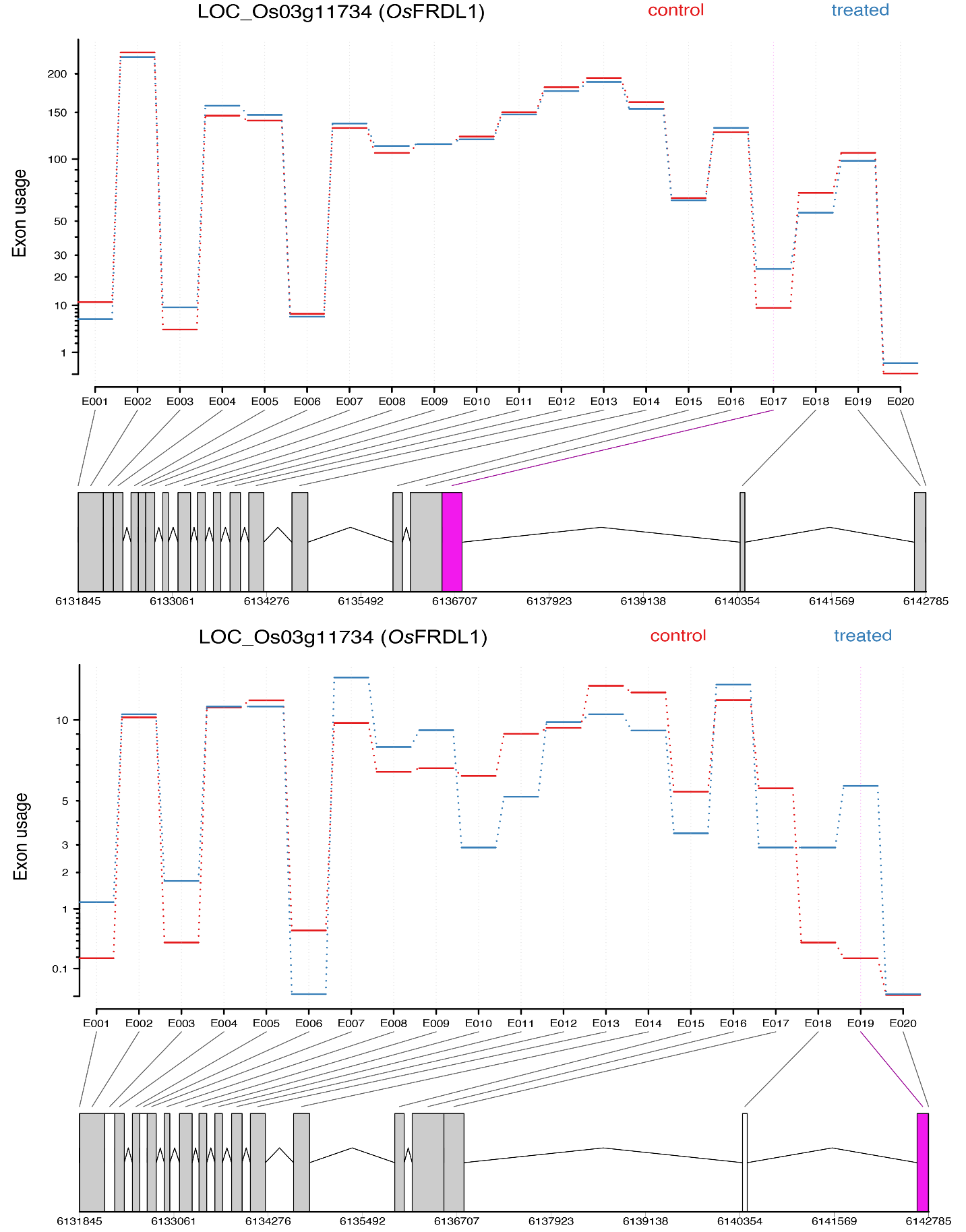

Figure S8: Representative figure showing the differential exon usage of OsFRDL1. Significant differential exons usages are highlighted.
