## Supplementary Table S2 for "Transcriptomic analysis revealed reactive oxygen species scavenging mechanisms associated with ferrous iron toxicity in aromatic Keteki Joha rice"

Supplementary Table S1: RNA-Seq samples showing the number of reads and percentage of uniquely mapped reads into the rice genome.

| **Tissue Type** | **Sample Code** | **No. of Reads**  **(Millions)** | **Uniquely Mapped**  **(%)** | **Condition** |
| --- | --- | --- | --- | --- |
| Root seedling | 1R1 | 10.10 | 72.27 | Untreated |
| Root seedling | 1R2 | 10.30 | 69.88 | Untreated |
| Root seedling | 1R3 | 10.60 | 70.39 | Untreated |
| Root seedling | 3R1 | 12.15 | 53.42 | Treated |
| Root seedling | 3R2 | 12.47 | 49.06 | Treated |
| Root seedling | 3R3 | 09.36 | 56.38 | Treated |
| Leaves seedling | 1S1 | 33.08 | 42.22 | Untreated |
| Leaves seedling | 1S2 | 37.40 | 41.59 | Untreated |
| Leaves seedling | 1S3 | 33.62 | 42.73 | Untreated |
| Leaves seedling | 3S1 | 26.88 | 44.31 | Treated |
| Leaves seedling | 3S2 | 26.09 | 49.24 | Treated |
| Leaves seedling | 3S3 | 31.25 | 47.58 | Treated |
| Leaves mature | 2S1 | 16.53 | 40.31 | Untreated |
| Leaves mature | 2S2 | 14.76 | 58.62 | Untreated |
| Leaves mature | 2S3 | 16.27 | 57.48 | Untreated |
| Leaves mature | 4S1 | 22.70 | 44.48 | Treated |
| Leaves mature | 4S2 | 24.36 | 46.59 | Treated |
| Leaves mature | 4S3 | 24.33 | 41.78 | Treated |
